# Post-transcriptional regulation of meiotic transcripts by the RNA binding protein CDM1 is associated with cytoplasmic condensate assemblies

**DOI:** 10.1101/2025.08.17.670746

**Authors:** Surendra Saddala, Neha Shukla, Jaroslav Fulnecek, Albert Cairo, Jakub Brolik, Pavlina Mikulkova, Jana Pecinkova, Anna Vargova, Sona Bukovcakova Valuchova, Claudio Capitao, Terezie Mandakova, Karel Riha

## Abstract

The transition from diploid to haploid life phases requires extensive reprogramming of gene expression that drive meiotic cell division and the differentiation of haploid forms. A hallmark of meiosis is widespread post-transcriptional regulation, including delayed translation that ensures timely protein production at specific stages. Here, we identify a mechanism controlling the translation of meiotic transcripts in Arabidopsis pollen mother cells, mediated by CALLOSE DEFECTIVE MICROSPORES1 (CDM1). We show that CDM1 affects several meiotic processes, including chromosome pairing, condensation, and cytokinesis. CDM1 forms dynamic, meiosis specific cytoplasmic foci that associate with components of processing bodies and stress granules, giving rise to three-phase condensate assemblies. Biochemical and cellular analyses reveal that CDM1 is an RNA-binding protein with an intrinsic ability to form ribonucleoprotein condensates. Furthermore, we demonstrate that CDM1 binds mRNAs expressed in prophase I and represses their translation until later meiotic stages, coinciding with the disassembly of CDM1 condensates. These findings establish CDM1 as a key post-transcriptional regulator that fine-tunes the expression of meiotic genes to ensure proper progression of microsporogenesis.

## Introduction

Meiosis is a specialized cellular process that enables the transition from diploid to haploid life forms through reductional cell division. During meiosis, a single diploid cell undergoes two consecutive rounds of chromosome segregation, producing four haploid cells. Depending on the organism’s sexual life cycle, these haploid cells may differentiate into gametes or develop into diverse multicellular structures. Thus, meiosis represents an important life cycle transition, marked by extensive changes in gene expression that drive both the execution of the meiotic program and the subsequent differentiation of haploid forms.

Compared to mitosis, which typically takes only tens of minutes, meiosis is a considerably slower process, often extending over several days ^1-4^. In some organisms, meiosis can pause and resume after prolonged periods, lasting months in the pollen mother cells of larch or even years in the case of human oocytes ^5, 6^. Structural changes in meiotic chromosome organization during the extended prophase I partially restrict transcription ^7, 8^, a limitation that is compensated by post-transcriptional mechanisms, which play a key role in fine-tuning the expression of specific transcripts required for meiotic progression and post-meiotic development ^9-11^.

Ribosome profiling in budding yeast has revealed that translation of many transcripts produced during prophase I is delayed until later meiotic stages ^12^. A prominent example is Clb3, a meiosis II-specific cyclin whose mRNA is translationally repressed during earlier stages of meiosis by the RNA-binding protein Rim4 ^13^. This repression is mediated through the sequestration of transcripts into amyloid-like aggregates, preventing premature translation ^14, 15^. The stabilization of meiotically produced mRNAs and their delayed translation is also hallmark of meiosis in mammals. For instance, during spermatogenesis, many genes involved in late meiotic processes are transcribed at early meiotic prophase, and mRNAs required for post-meiotic sperm differentiation are produced already during pachytene ^16^. Post-transcriptional gene regulation is equally important for development of oocytes. Mammalian oocytes are transcriptionally inactive and the completion of meiosis as well as early embryonic development relies on maternally synthesized mRNAs ^17^. These mRNAs are stored in mitochondria-associated ribonucleoprotein compartments formed by the RNA-binding protein ZAR1 ^18^.

Post-transcriptional regulation of meiotic gene expression is much less explored in plants, yet emerging evidence highlight its critical role in plant germline differentiation. In European larch, male meiosis enters a prolonged arrest during diplotene over winter, characterized by periodic chromatin relaxation accompanied by episodic transcriptional activity ^5^. These transcripts are not fully spliced, but retained in nuclear Cajal bodies, preventing their premature translation until meiosis resumes ^19^. This mechanism underscores how post-transcriptional control ensures temporal precision in gene expression, aligning meiotic progression with seasonal changes.

Post-transcriptional regulation also governs the exit from meiosis and transition to post-meiotic development in Arabidopsis. This transition is regulated by meiotic (M) bodies, cytoplasmic condensates composed of a processing (P) body core surrounded by a stress granule (SG)-like shell ^20^. At the end of meiosis, M-bodies transiently sequester translation initiation factor eIFiso4G2 via the meiosis-specific protein TDM1, thereby temporarily repressing translation to facilitate termination of the meiotic program ^21^. TDM1 is recruited to M-bodies by SMG7, an evolutionary conserved nonsense-mediated RNA decay factor that acts as an adaptor protein involved in P-body remodeling and regulation of its size homeostasis ^20^. The SMG7-TDM1 module is negatively regulated by meiosis I specific cyclin TAM, ensuring that meiotic exit does not occur prematurely during interkinesis ^22-24^.

Allele-specific transcriptomics in maize revealed that transcripts generated in diploid pollen precursor cells persist in haploid microspores for up to 11 days post-meiosis ^25^, suggesting a strategy of mRNA storage and delayed processing that supports post-meiotic development. However, the underlying molecular mechanisms remain unknown. In this study, we describe a mechanism that delays the expression of meiotically produced transcripts during *Arabidopsis* microsporogenesis, mediated by CALLOSE DEFECTIVE MICROSPORE1 (CDM1; AGI gene code At1G68200; also known under alternative name C3H15). CDM1 belongs to the family of tandem CCCH-type zinc-finger proteins ^26^. Previous work has shown that CDM1 affects callose metabolism during microsporogenesis, and that its inactivation leads to abortive microspore development and male sterility ^27, 28^.

Here, we show that CDM1’s function extends beyond callose metabolism, influencing multiple meiotic processes including chromosome pairing, condensation, and cytokinesis. CDM1 localizes to dynamic cytoplasmic speckles that are prominent during prophase I. These speckles closely associate with M-bodies, forming three-phase condensate assemblies, suggesting a coordinated role in post-transcriptional regulation during meiosis. We further demonstrate that CDM1 binds mRNAs transcribed during prophase I and represses their translation until subsequent meiotic stages. These results indicate that CDM1 acts as a post-transcriptional regulator, fine-tuning the translation of meiotically expressed genes to ensure proper progression of microsporogenesis.

## Results

### CDM1 deficiency affects multiple meiotic processes

To identify regulators of meiotic progression in Arabidopsis, we performed a suppressor genetic screen in *Arabidopsis smg7-6* mutants, which are deficient in meiotic exit and exhibit a tenfold decrease in viable pollen along with reduced fertility ^29, 30^. Screening an ethyl methanesulfonate (EMS)-mutagenized *smg7-6* population for increased seed yield revealed a line that restored the viable pollen count to wild-type levels (Fig. S1). Further genetic analyses and association mapping identified a recessive mutation in the CDM1 gene as the most likely causative variant (Fig. S1; Table S1). This mutation results in a substitution of proline with serine at position 152. Plants carrying the homozygous mutant allele, designated *cdm1-2*, do not exhibit any apparent phenotype aside from rescuing meiotic defects in the *smg7-6* background (Fig. S1).

Identification of CDM1 in the suppressor screen prompt us to analyze its meiotic role in more detail. A previous study characterized a T-DNA insertion allele (*cdm1-1*) that disrupts the CDM1 gene between the C-terminal zinc-finger motifs ^27^. While that study reported reduced pollen count as well as aberrant callose deposition and dissolution during the tetrad stage, meiosis was not characterized. Our analysis of *cdm1-1* mutants revealed a drastic reduction in the number of viable pollen, some of which appeared larger compared to the wild type (Fig. 1a,b). The mutants were almost completely infertile, and the few viable seeds that were recovered were also substantially larger (Fig. 1c). Callose staining revealed multiple cytokinetic defects. First, we observed initiation of premature cytokinesis during meiotic interphase that was not fully completed (Fig. 1d). This was confirmed by analysis using a plasma membrane marker line (*GFP-SYP132*), which showed plasma membrane invaginations at the equatorial plate during interkinesis but no full separation of the segregated nuclei (Fig. S2). Second, the callose cell wall was thinner at the tetrad stage, confirming its aberrant deposition ^27^. Finally, we observed that some haploid nuclei at the tetrad stage were not separated by a cell wall, resulting in binuclear tetrad cells (Fig. 1d).

**Figure 1.**
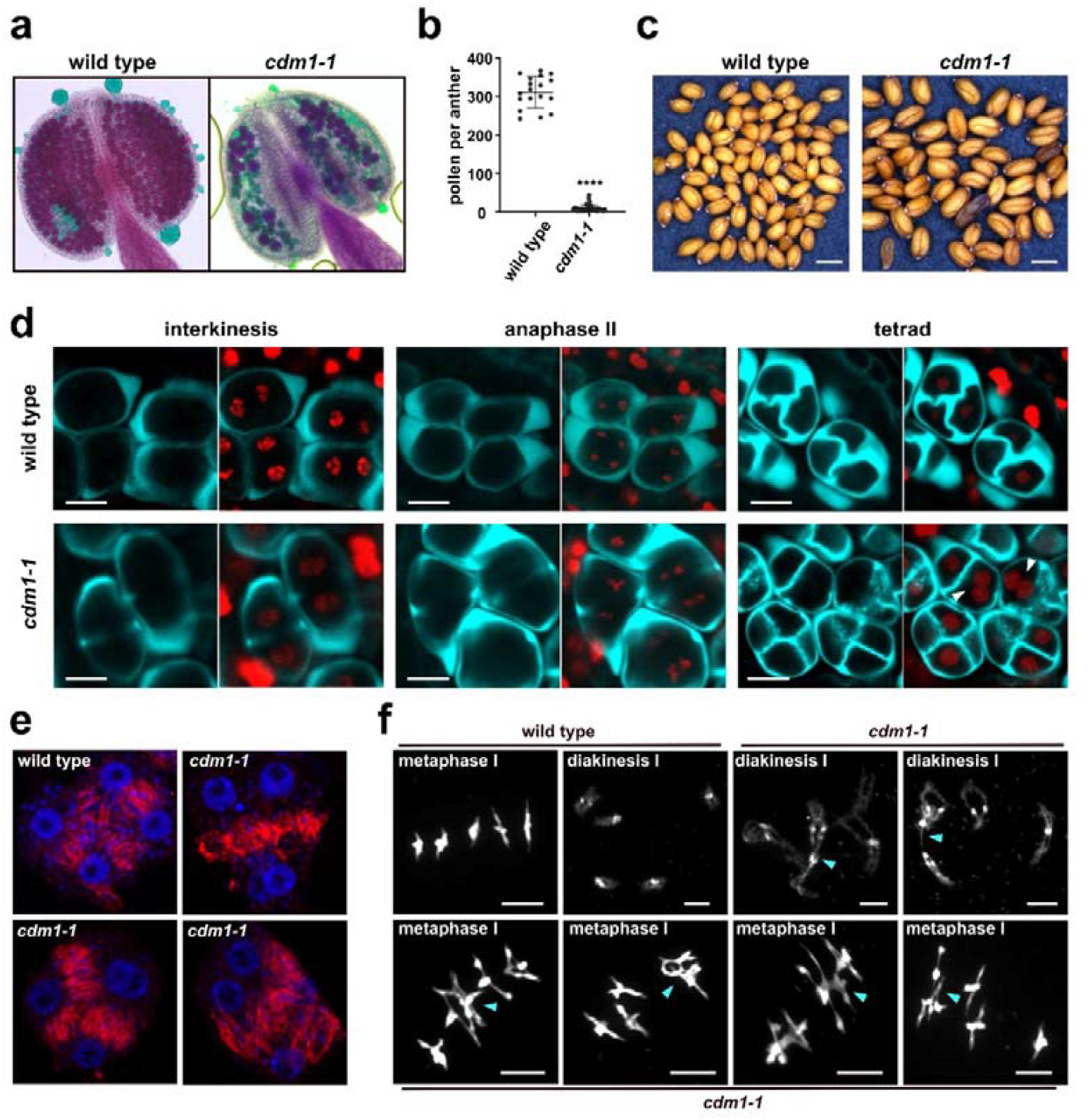
Meiotic and post-meiotic defects in *cdm1-1* mutants. **(a)** Alexander staining of wild type and *cdm1-1* mutant anthers. **(b)** Quantification of pollen viability shown in (a) (unpaired t-test, *p* < 0.0001). **(c)** Seeds from wild type and *cdm1-1* mutants (scale = 0.5 mm). (**d**) SR2200 staining of callose (cyan) in meiocytes expressing the HTA10-RFP chromatin marker (red), imaged by confocal microscopy (scale = 10 µm). (**e**) Anti-tubulin immunostaining (red) during telophase II showing radial microtubule arrays (RMAs). DNA is counterstained with DAPI (blue). (**f**) Meiotic chromosome spreads stained with DAPI. Association of heterologous bivalents are indicated by blue arrowheads.

In Arabidopsis male meiosis, cytokinesis takes place simultaneously after completion of both nuclear divisions and is characterized by centripetal ingrowth of the primary cell wall. The position of the future cell wall is determined by radial microtubule arrays (RMAs), which interconnect all haploid nuclei ^31^. In certain *cdm1-1* meiocytes, however, RMAs were observed forming between only two or three nuclei (Fig. 1e), while being absent between others, which could account for the formation of binuclear tetrad cells (Fig. 1d).

To investigate whether CDM1 influences meiotic processes beyond cytokinesis, we performed chromosome spread analyses in *cdm1-1* mutants. At diakinesis, mutant chromosomes appeared less condensed compared to wild type controls and frequently exhibited associations between heterologous bivalents (Fig. 1f). These aberrant inter-bivalent interactions persisted into metaphase I, where chromosomes were fully condensed (Fig. 1f), suggesting potential ectopic recombination or mispairing between non-homologous chromosomes. Together, this detailed phenotypic analysis of *cdm1-1* mutants demonstrates that the function of CDM1 is not solely involved in callose metabolism, as previously reported ^27^, but influences multiple meiotic processes, including chromosome organization during prophase I.

### CDM1 forms dynamic cytoplasmic speckles associated with M-bodies

To further investigate the meiotic function of CDM1, we performed expression and intracellular localization studies. We generated C-terminal fusions of the CDM1 gene with endogenous promoter, to GUS, GFP, and TagRFP reporters. All reporter constructs restored fertility in *cdm1-1* mutants, confirming their functionality (Fig. S3). The *CDM1:GUS* construct showed patchy expression in the vasculature of roots of young seedlings, as well as strong staining in anthers of floral buds at the meiotic stage (Fig. S4). The staining disappeared in older floral buds and open flowers, indicating that CDM1 expression is restricted to meiosis. This expression pattern is consistent with previous observations obtained using *pCDM1::GUS* promoter fusion constructs ^28^.

To examine CDM1 subcellular dynamics, we performed live imaging of pollen mother cells in CDM1-GFP plants (Video 1). The GFP signal appeared at early prophase I, forming cytoplasmic foci that were most prominent during mid-prophase I and partially dissolved following nuclear envelope breakdown at the onset of chromosome segregation. The signal was no longer detectable by the end of meiosis. To gain more detailed insights, we conducted confocal microscopy on fixed pollen mother cells with chromosomes counterstained with DAPI (Fig. 2a). This analysis confirmed prominent speckles during zygotene and pachytene, with both their number and size decreasing as cells progressed through metaphase I into meiosis II (Fig. 2b,c). Although some foci persisted in later stages, the CDM1 partition coefficient, which measures the signal ratio between foci and cytoplasm, declined (Fig. 2d), consistent with the partial dissolution observed in live imaging (Video 1). Notably, a portion of CDM1 appeared to associate with the spindle during meiosis I and II (Fig. 2a). Collectively, these data demonstrate that CDM1 exhibits a highly dynamic localization pattern, forming prominent cytoplasmic foci during prophase I and redistributing to the cytoplasm and spindle during chromosome segregation. Although *cdm1-1* mutants are female fertile ^27^, CDM1 is also expressed in megaspore mother cells, where it forms similar cytoplasmic foci (Fig. S5).

**Figure 2.**
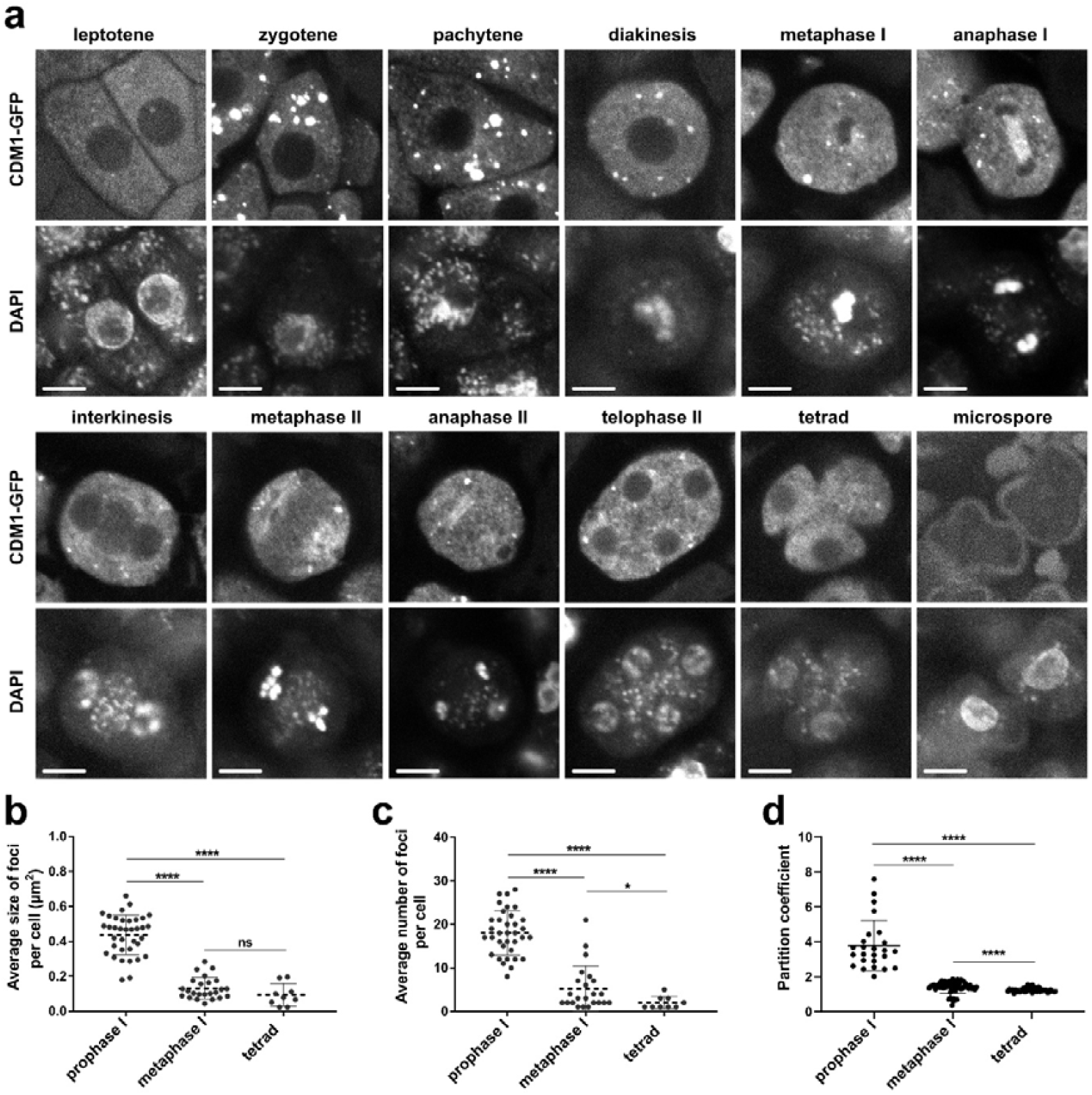
Subcellular localization of CDM1-GFP during male meiosis. (**a**) Confocal micrographs of pollen mother cells expressing CDM1-GFP (scale = 5 µm). (**b-d**) Dot plots showing quantification of the average size of CDM1-GFP foci per cell (b), average number of CDM1-GFP foci per cell (c), and partitioning coefficient of individual foci (d) at indicated meiotic stages. Data were quantified from a single representative confocal layer. Statistical significance of the difference was assessed using unpaired Mann-Whitney test. *P*-values: **** = <0.0001, * = 0.0250, ns = not significant.

We next assessed whether CDM1 speckles associate with M-bodies. To this end, we performed pairwise co-localization of CDM1 with P-body markers (SMG7-TagRFP and DCP1-GFP), which label the M-body core, and with the SG marker TagRFP-RBP47, which localizes to the surrounding shell ^20^. Confocal microscopy in pachytene meiocytes revealed extensive co-localization of CDM1 foci with all three markers, indicating a close association with M-bodies (Fig. 3a–i). However, closer inspection suggested that CDM1 signals were juxtaposed rather than fully overlapping with the P-body markers (Fig. 3b,e). In contrast, overlap with TagRFP-RBP47 was more pronounced (Fig. 3h), consistent with CDM1 localization to the M-body shell.

**Figure 3.**
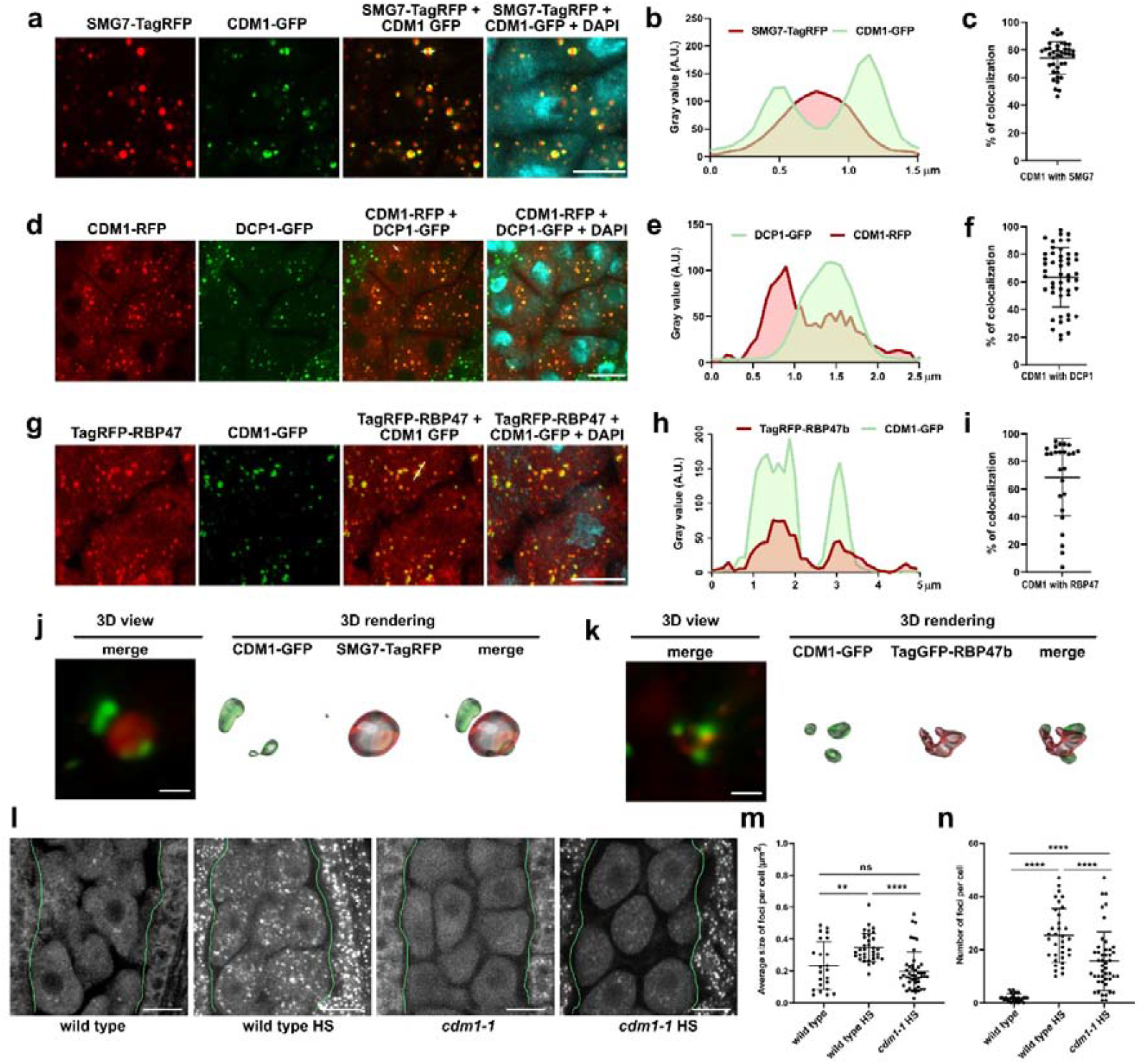
Colocalization of CDM1 with markers of M-bodies. (**a**) Confocal micrographs of zygotene meiocytes co-expressing CDM1-GFP and SMG7-TagRFP (scale bar = 10 µm). (**b**) Diagram showing superimposed intensity profiles of GFP and TagRFP signals along the line indicating in micrograph (a). (**c**) Dot plot indicating precenting of CDM1 foci co-localizing with SMG7 foci. (**d**) Confocal micrographs of zygotene meiocytes co-expressing CDM1-RFP and GFP (scale bar = 10 µm). (**e**) Diagram showing superimposed intensity profiles of RFP and GFP signals along the line indicating in micrograph (d). (**f**) Dot plot indicating precenting of CDM1 foci co-localizing with DCP1 foci. (**g**) Confocal micrographs of zygotene meiocytes co-expressing CDM1-GFP and TagRFP-RBP47 (scale bar = 10 µm). (**h**) Diagram showing superimposed intensity profiles of GFP and TagRFP signals along the line indicating in micrograph (g). (**i**) Dot plot indicating precenting of CDM1 foci co-localizing with RBP47 foci. (**j**,**k**) Supper-resolution micrographs of indicated protein condensates visualized by 3D view and 3D rendering using Imaris software (scale bar = 0.5 µm). (**l**) Confocal micrographs of wild type and cdm1-1 meiocytes expressing TagRFP-RBP47b construct. Heat shock (HS) treated meiocytes were exposed to 39°C for 20 min. Green lines separatee pollen mother cells from tapetal cells (scale bar = 10 µm). (**m**,**n**) Dot plots showing quantification of the average size of TagRFP-RBP47b foci per cell (m), and average number of the foci per cell (n). Statistical significance of the difference was assessed using unpaired Mann-Whitney test. *P*-values: **** = <0.0001, ** = 0.004, ns = not significant.

To refine these observations, we used structured illumination-based super-resolution microscopy. This analysis confirmed that CDM1 and SMG7 occupy adjacent, non-overlapping domains, with CDM1 foci encircling the SMG7-labeled core (Fig. 3j). Surprisingly, CDM1 did not fully co-localize with RBP47b either (Fig. 3k). Three-dimensional rendering revealed that CDM1 and RBP47b form distinct but intermingled patches surrounding the presumed M-body core (Fig. 3k). Together, these results suggest that pachytene M-bodies are three-phase condensates, composed of a P-body core surrounded by discrete, interspersed SG-like and CDM1 condensates.

Spontaneous formation of SG-like foci, marked here by RBP47b, is under normal conditions specific to meiocytes and does not occur in the surrounding somatic cells ^20^ (Fig. 3l). Therefore, we asked whether CDM1, which is exclusively expressed in meiocytes, contributes to the nucleation of these foci. Indeed, confocal microscopy revealed that RBP47b foci are completely abolished in *cdm1-1* mutants (Fig. 3l). Heat stress induces the condensation of SGs in somatic cells and leads to their pronounced increase in meiocytes ^20^ (Fig. 3l-n). Strikingly, heat-induced SGs form much less efficiently in CDM1-deficient meiocytes than in the surrounding somatic cell (Fig. 3l-n). This suggests that CDM1 not only nucleates the SG-like shell of M-bodies under normal conditions but also facilitates the condensation of canonical SGs in meiocytes under heat stress.

### CDM1 represses translation of late meiotic proteins during prophase I

The pleiotropic effects of CDM1 on meiosis, its localization to cytoplasmic speckles, and the previously reported RNA-binding capacity of the closely related Arabidopsis protein C3H14 ^32^ suggested that CDM1 may bind RNA and regulate meiotically expressed transcripts. To identify potential targets of CDM1 regulation, we performed CDM1:GFP immunoprecipitation from meiotic floral buds using anti-GFP nanobeads, followed by sequencing of the co-purified RNA. Genes enriched in the CDM1:GFP RIP-seq compared with the wild-type RIP-seq control (Table S2) were further filtered based on altered expression in *cdm1-1* mutants in microarray data ^27^ and specific expression in floral buds, according to the Arabidopsis transcriptome database (https://travadb.org/) ^33^. Two preselected candidates, *G3BP5* (*At3g07250*) and *BT3* (*At1g05690*), were validated by RIP-qPCR for association with CDM1:GFP (Fig. 4a) and by RT-qPCR as being downregulated in *cdm1-1* mutants (Fig. 4b) and were analyzed further.

**Figure 4.**
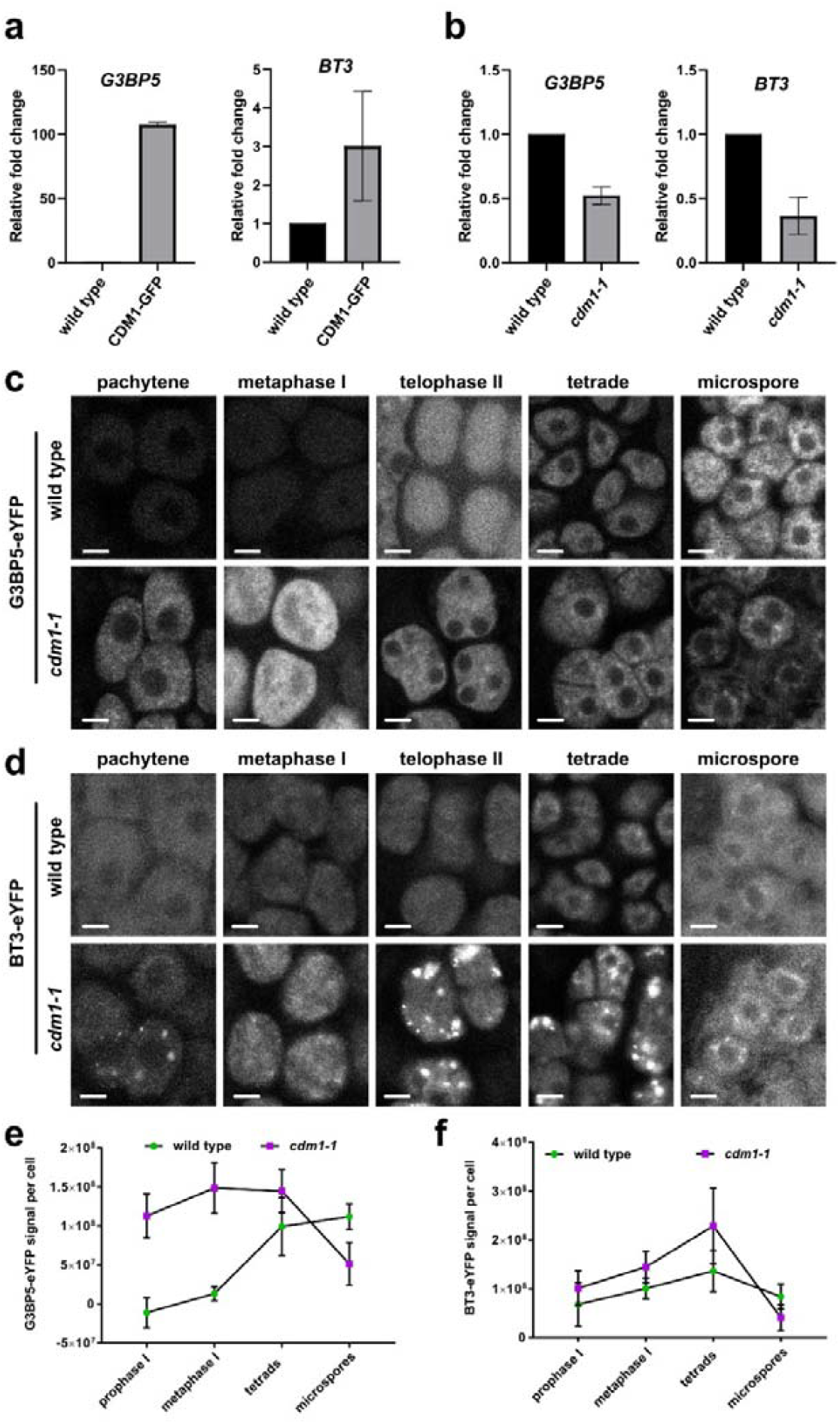
Regulation of *G3BP5* and *BT3* in pollen mother cells by CDM1. (**a**) Bar charts showing enrichment of the indicated mRNAs in anti-GFP immunoprecipitated fractions from wild type and CDM1-GFP plants, as measured by RIP-qPCR. Error bars indicated standard deviations from two biological replicates. (**b**) Relative abundance of *G3BP5* and *BT3* mRNAs in floral buds from wild type and *cdm1-1* plants quantified by RT-qPCR. Error bars indicate SD from three biological replicates. (**c**,**d**) Confocal micrographs of G3BP5-YFP (c) and BT3-YFP (d) signals in wild type and *cdm1-1* pollen mother cells. All images in (c) and (d) were acquired using identical microscope settings and processed with the same parameters. Images show single representative confocal sections; scale bars = 5 µm. (**e, f**) Quantification of G3BP5-YFP (e) and BT3-YFP (f) total fluorescence intensity per meiocyte.

To monitor G3BP5 and BT3 protein expression during meiosis, we generated genic constructs containing native promoter regions fused to eYFP at the C-terminus. In wild type, G3BP5:eYFP showed little to no signal during meiosis I, with expression rising in tetrads and peaking in microspores (Fig. 4c). BT3:eYFP displayed gradual increase through meiosis I with the highest signal in tetrads (Fig. 4d). In *cdm1-1*, however, both proteins accumulated earlier and at higher levels. G3BP5:eYFP was already prominent at pachytene, reached a maximum at metaphase I, and declined by the microspore stage (Fig. 4c,e). BT3:eYFP in *cdm1-1* formed distinct cytoplasmic foci, first visible at pachytene, with peak intensity at telophase II and the tetrad stage, followed by a sharp decrease in microspores (Fig. 4d,f). These foci, absent in wild-type meiocytes, may represent aggregates of overproduced protein.

To determine whether CDM1 regulates *G3BP5* and *BT3* expression at the transcriptional or post-transcriptional level, we performed single-molecule RNA FISH in pachytene meiocytes. For *G3BP5*, we detected an average of ∼400 foci per cell, with ∼300 in the cytoplasm and ∼100 in the nucleus (Fig. 5a,b). In *cdm1-1* mutants, cytoplasmic foci were significantly reduced, whereas nuclear foci remained unchanged. A similar pattern was observed for *BT3* transcripts, which were less abundant than *G3BP5* mRNAs but likewise showed a marked reduction in the absence of CDM1 (Fig. 5c,d). The RNA-FISH data are consistent with the results of RT-qPCR (Fig.4b). These findings indicate that *G3BP5* and *BT3* mRNAs are already present during pachytene, where they are translationally repressed and likely stabilized by CDM1. Loss of CDM1 appears to relieve this repression, resulting in strong protein accumulation (Fig. 4c–f). This premature translation may deplete the mRNA pool during prophase I, consistent with the reduced transcript numbers (Fig. 5) and lower protein levels at the end of meiosis (Fig. 4c–f). Thus, CDM1 functions as a translational repressor, delaying expression of late meiotic and post-meiotic proteins from mRNAs synthesized during early meiotic stages.

**Figure 5.**
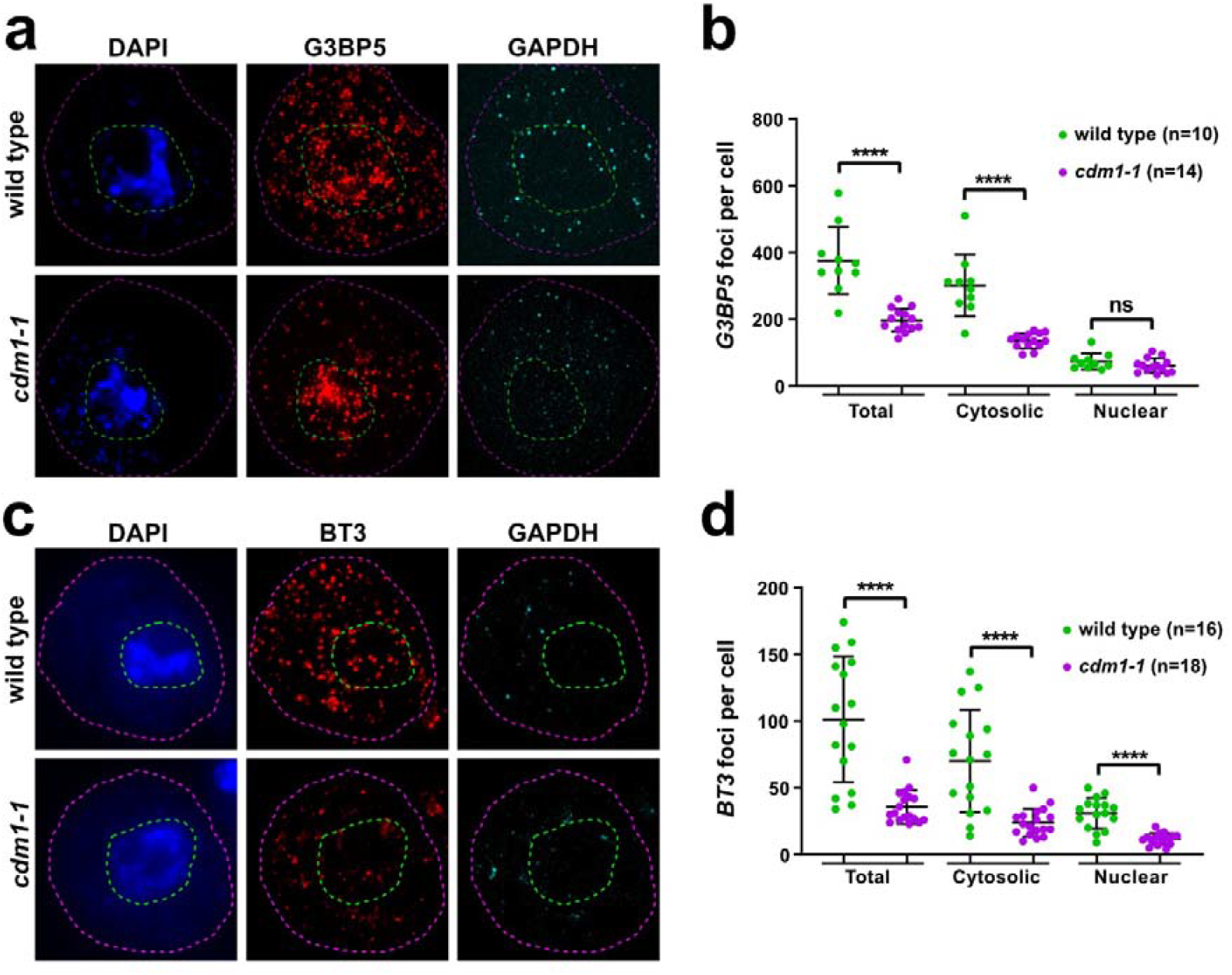
smFISH analysis of *G3BP5* and *BT3* transcripts in wild-type and *cdm1-1* meiocytes. (**a,c**) Micrographs of pachytene meiociytes subject to smFISH with G3BP5 (a) or BT3 (c) probes (red). DNA was counterstained with DAPI (blue), and a probe detecting ubiquitously expressed GAPDH was used as a control (cyan). Magenta dotted lines outline cell boundaries; green dotted lines indicate approximate nuclear boundaries inferred from the DAPI signal. (**b, d**) Quantification of mRNA foci per cell for *G3BP5* (b) and *BT3* (d). Statistical significance was assessed using an unpaired Mann-Whitney test (**** = <0.0001, ns = not significant). Scale bar = 5 µm.

### CDM1 is an RNA binding protein with propensity to form molecular biocondensates

The role of CDM1 in translational repression suggests that it may directly bind to its target mRNAs. To test this, we performed RNA electrophoretic mobility shift assays (REMSA) using *E. coli* - expressed CDM1 and oligoribonucleotides spanning 5⍰UTR of *G3BP5* mRNA, as UTRs are known as interaction hubs for translational regulators ^34^. REMSA with three oligoribonucleotides spanning different regions of the 5⍰UTR revealed a CDM1-dependent mobility shift with probe 5⍰-1, corresponding to the 77 nt at the 5⍰ terminus (Fig. S6 and Fig. 6a). Competition assay with excess unlabeled RNAs confirmed the high specificity of this interaction (Fig. 6a), demonstrating that CDM1 can specifically bind the 5⍰UTR of its target RNA.

**Figure 6.**
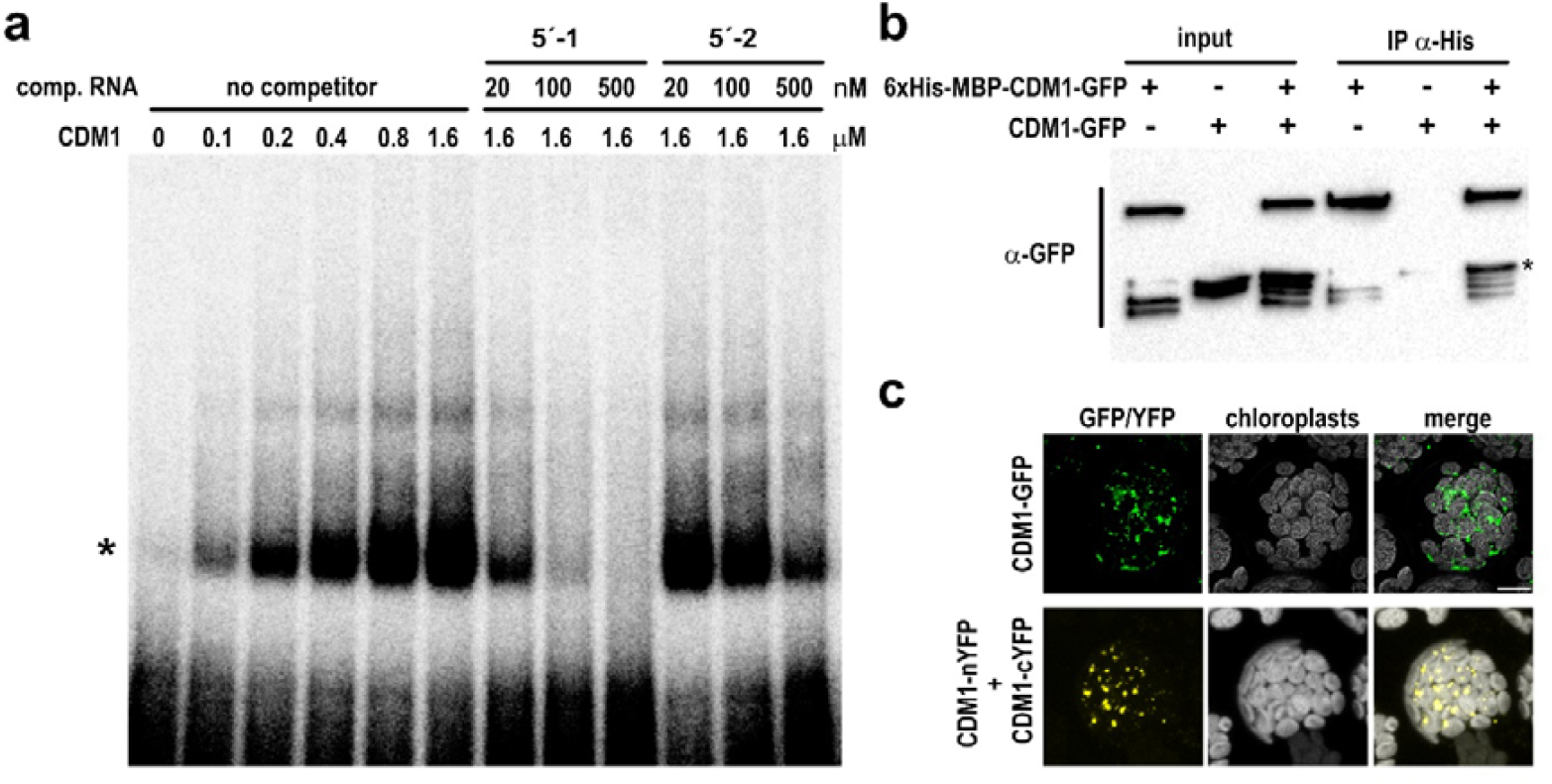
CDM1 directly binds *G3BP5* RNA and self-associates. (**a**) Autoradiogram of REMSA with CDM1 protein and a 20 nM 5′-1 RNA probe spanning the first 75 nucleotides of *G3BP5*′ 5′UTR. Unlabeled 5′-1 and 5′-2 RNAs were used as competitors. The asterisk indicates the CDM1-GRBP5 complex (**b**). Western blot of *in vitro* pull-down assay with His-MBP-CDM1-GFP and CDM1-GFP proteins using anti-His antibody. Proteins were detected using anti-GFP antibody. The asterisk marks the CDM1-GFP signal. (**c**) Bimolecular fluorescence complementation assay in mesophyll protoplasts. Top panel: micrographs showing CDM1-GFP expression in transfected protoplasts. Bottom panel: YFP fluorescence signal upon co-transfection of CDM1 constructs fused to either N-terminal (CDM1-nYFP) or C-termina (CDM1-cYFP) halves of YFP.

We noticed that heterologously expressed CDM1 tends to aggregate *in vitro*. The localization of CDM1 to cytoplasmic speckles and its tendency to aggregate prompted us to test whether it can homomultimerize. *In vitro* immunoprecipitation using soluble MBP-CDM1 fusion proteins revealed that CDM1 forms dimers or higher-order multimers (Fig. 6b), and a specific self-interaction was also confirmed by yeast two-hybrid assay (Fig. S7). To assess interactions in plant cells, we performed bimolecular fluorescence complementation (BiFC) in mesophyll protoplasts. Ectopically expressed CDM1-GFP formed prominent cytoplasmic speckles resembling those in pollen mother cells (Fig. 6c). Co-transfection with bimolecular complementation constructs produced similar foci, indicating direct interaction or close spatial association of CDM1 molecules within these structures.

Close association of CDM1 speckles with P-bodies and SGs in the form of multiphasic M-bodies, suggests that CDM1 can assemble into biocondensates with properties of liquid-liquid phase-separated droplets. One hallmark of such droplets is their ability to fuse, a property observed for CDM1 speckles both in meiocytes and in heterologously expressed CDM1 speckles in mesophyll protoplasts (Fig. 7a). Another defining feature is the high mobility of their constituents. Fluorescence recovery after photobleaching (FRAP) experiments showed partial restoration of the CDM1-GFP signal, indicating exchange of CDM1 molecules with the surrounding cytoplasm (Fig. 7b,c).

**Figure 7.**
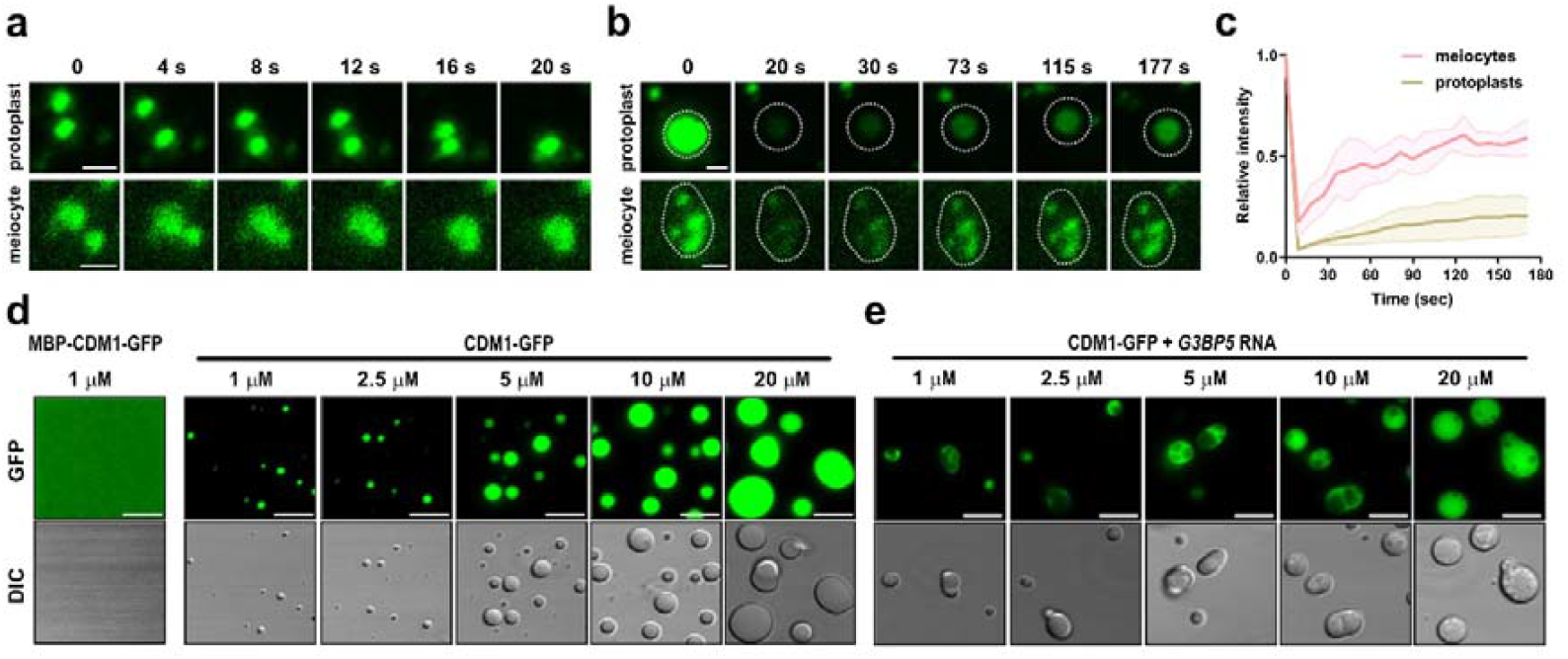
Dynamic properties of CDM1 foci and *in vitro* phase separation of CDM1 protein. (**a**) Time-lapse series showing fusion of two CDM1-GFP foci in protoplasts (top) and meiocytes (bottom). (**b**) FRAP analysis of CDM1-GFP signal in protoplasts (top) and in meiocytes (bottom). Scale bars in (a) and (b) = 1 µm. (**c**) Quantification of CDM1-GFP FRAP in meiocytes and protoplasts. (**d**,**e**) *In vitro* phase separation assay with increasing concentration of CDM1-GFP, either without (d) or with (e) *G3BP5* RNA (100 ng/µL). Top panels show GFP signal and the bottom panels show the corresponding DIC images. Scale bar = 5 µm.

To determine whether CDM1 possesses intrinsic phase-separation capabilities, we performed *in vitro* droplet formation assays using MBP-CDM1-GFP fusion protein. Cleavage of the MBP solubility tag induced the formation of liquid droplets, whose size increased with protein concentration (Fig. 7d). To test the influence of RNA, we repeated the assay in the presence of a 5′-end fragment of the *G3BP5* RNA. RNA facilitated CDM1 phase separation, producing larger droplets at 1 µM and 2.5 µM concentrations compared with corresponding controls lacking RNA (Fig. 7d,e). Notably, CDM1 formed biphasic condensates in the presence of RNA (Fig. 7e), suggesting the existence of two demixed populations, potentially representing RNA-bound and RNA-free configurations.

Together, these results demonstrate that CDM1 speckles in plant cells exhibit characteristics of liquid-liquid phase-separated droplets, driven by an intrinsic ability of CDM1 to condense such structures, particularly in the presence of RNA.

## Discussion

Post-transcriptional gene regulation plays a central role in meiosis and germline differentiation. In yeasts and animals, many components of the regulatory networks governing mRNA processing, translation, and storage are concentrated in germline-specific RNP assemblies. These condensates regulate precise timing of protein expression at the appropriate developmental stage, safeguard germline identity by repressing somatic gene expression, and protect genome integrity by counteracting transposon activation ^14, 18, 35-40^.

Post-transcriptional mechanisms also play an important role in the differentiation of the plant germline, as evidenced mainly by studies on phasiRNA- and microRNA-mediated regulation ^41, 42^. However, little is known about the mechanisms that control mRNA storage and the timing of its translation. Here, we identify CDM1 as a key regulator of translation in *Arabidopsis* pollen mother cells. We show that CDM1 is an RNA-binding protein specifically expressed in meiotic cells, where it controls the translation of its target mRNAs. CDM1/C3H15 belongs to the tandem CCCH zinc finger protein family, whose members are implicated in diverse biological processes, including hormone signaling, seed germination, stress responses, and pathogen defense ^43^. Although the molecular functions of most of these proteins remain to be elucidated, several of them, including C3H14, the closest homologue of CDM1, have been found to associate with stress granules and P-bodies or bind to RNA, suggesting their role in RNA metabolism ^26, 32, 44-47^. Indeed, AtTZF1, an archetypal member of this family, was recently shown to regulate salt stress tolerance by promoting decay of mRNA encoding a tonoplast-localized calcium pump ^48^.

A recent study using overexpression lines suggested that CDM1/C3H15 acts as a transcriptional repressor of *HEAT SHOCK TRANSCRIPTION FACTOR A2* ^49^. However, transcriptional regulation is unlikely to be the primary mechanism of CDM1 action in Arabidopsis meiosis. Our analysis of functionally complementing reporter lines expressing CDM1:GFP/RFP fusion proteins under the control of the native promoter did not reveal any prominent nuclear localization at any stage of meiosis (Fig. 2). Furthermore, CDM1-mediated repression of G3BP5 and BT3 in prophase I appears to occur at the level of protein synthesis rather than mRNA production (Fig. 4 and 5). Finally, smFISH analysis of *G3BP5* mRNA showed that CDM1 primarily affects the abundance of cytoplasmic transcripts, but does not seem to influence their nuclear pool (Fig. 6), suggesting a role in cytoplasmic mRNA stabilization rather than transcription.

These data indicate that CDM1 acts as a translational repressor of mRNAs synthesized early in meiosis and delays their expression until late meiotic or post-meiotic stages. Indeed, while the function of G3BP5 is unknown, BT3 was implicated in post-meiotic pollen development ^50^. Notably, translational repression coincides with the formation of prominent cytoplasmic CDM1-foci in prophase I. By analogy to stress granules, CDM1 may sequester specific meiotic transcripts within cytoplasmic biocondensates, thereby inhibiting their translation and protecting them from degradation. In the absence of CDM1, G3BP5 and BT3 are prematurely translated in prophase I, and their mRNA levels decline, presumably through co-translational RNA decay ^51-53^. Partial dissolution of CDM1 granules at the onset of meiotic divisions correlates with the accumulation of G3BP5 and BT3 proteins, suggesting release of their transcripts from translational repression. CDM1 does not fully disappear during the divisions. It persists in the cytoplasm and partially accumulates to meiotic spindles. Translation initiation factors such as eIFiso4G2 also localize to meiotic spindles, which suggests that these structures may serve as sites of enhanced translation ^21^. The role of CDM1 in translational regulation therefore appears more complex than acting solely as a repressor and it may shift with its ability to condense in the context of meiotic divisions.

Our previous studies highlighted the importance of cytoplasmic RNP condensates in the regulation of Arabidopsis meiosis. We characterized M-bodies as biphasic structures consisting of a P-body core and an SG-shell that co-ordinate the termination of meiosis and the transition to post-meiotic development ^20, 21^. Here we show that CDM1 is an essential component of M-bodies, forming three-phase condensate assemblies during prophase I. CDM1 foci surrounding the P-body core are crucial for seeding the SG-shell, although they appear to form spatially distinct domains. The ability of CDM1 to bind RNA and condense may locally enrich translationally inactive mRNAs and further nucleate the formation of SG-like condensates.

M-bodies are complex and highly dynamic structures changing their composition in the course of meiotic progression. While CDM1 and SG-components are most prominently present in prophase I, TDM1 and the eIFiso4G2 translation initiation factor associate with the P-body core during meiosis II ^20, 21^. Importantly, these composition shifts coincide with different functionalities of M-bodies during meiotic progression. In prophase I, CDM1 confers temporal translational repression and presumably storage of late meiotic or post-meiotic transcripts, whereas SMG7 and TDM1 appear to facilitate termination of meiosis by permanent inactivation of meiotic genes ^21^. Although CDM1 and SMG7-TDM1 form spatially distinct phases, genetic interaction between *CDM1* and *SMG7* (Fig. S1) indicates that organization of these assemblies into M-bodies is functionally meaningful and may facilitate orchestration of gene expression during plant germline differentiation. These findings parallel the complex behavior of cytoplasmic RNP condensates in animal oogenesis ^18, 40, 54^, suggesting that dynamic RNP compartmentalization represents an evolutionarily conserved mechanism underpinning germline development across kingdoms.

In conclusion, our findings establish CDM1 as a key component of the post-transcriptional machinery in Arabidopsis meiosis, acting as a temporal regulator of the translation of meiotic and post-meiotic transcripts. The dynamic integration of CDM1 into M-bodies highlights the functional versatility of these condensates as hubs of translational control, whose composition and activity alter during meiotic progression. A recent observation of temperature-induced recruitment of the meiotic A-type cyclin TAM into SGs and M-bodies, and the significance of this process for meiotic fidelity under heat stress ^55^, supports the idea that these RNP assemblies also modulate the meiotic program in response to environmental cues. Thus, M-bodies represent a unique system for studying multi-phase RNP biocondensates and their role in regulating the precise timing of gene expression during developmental transitions and stress responses.

## Materials and Methods

### Plant material and growth conditions

*Arabidopsis thaliana* ecotype Col-0, mutant lines *cdm1-1* (SALK_065040)^27^ and *smg7-6* ^30^, as well as reporter lines, were grown in soil under controlled long-day conditions (16⍰h light/ 8⍰h dark) at 21⍰°C and 50-60% relative humidity. The following Arabidopsis reporter lines were used in this study: *SMG7-TagRFP* ^21^, *DCP1-G3GFP* ^20^, *TagRFP-RBP47b* ^20^, *HTA:RFP* ^4^, and *GFP-SYP132* ^56^. The CDM1 reporter constructs were generated by amplifying the genomic sequence of *CDM1*, including 1500⍰bp upstream of the start codon but excluding the stop codon, using primers pCDM1-F and CDM1RSTOP-(Table S3). The PCR product was cloned into the pENTR™/D-TOPO™ vector (Invitrogen, cat. no. 45-0218) to generate an entry clone, which was subsequently recombined into pGWB633, pGWB650, and pGWB659 ^57^ by LR reactions (Invitrogen, cat. no. 11791-020), resulting in the constructs *CDM1:GUS, CDM1:G3GFP*, and *CDM1:TagRFP*, respectively. The *G3BP5:eYFP* and *BT3:eYFP* reporter constructs were generated analogously by amplifying genomic regions of these genes, including their promoters, using primers listed in Table S3. The PCR products were cloned into pENTR™/D-TOPO™ and recombined into pGWB640. All constructs were introduced into *cdm1-1*^*+/-*^ plants via *Agrobacterium tumefaciens-*mediated transformation using the floral dip method to establish corresponding reporter lines.

### Suppressor screen

The suppressor screen for mutations that restore fertility in *smg7-6* mutant plants was performed as previously described ^29^. Causal *de novo* mutations associated with the suppressor phenotype were identified by whole-genome sequencing and mapped using the ArtMAP pipeline ^58^. The *cdm1-2* mutation was genotyped using the High-Resolution Melting (HRM) analysis method with primers HRM_EMSCDM1105_F and HRM_EMSCDM1105_R (Table S3).

### Cytology

Staining of pollen mother cells in whole anthers was performed as previously described ^29^ with 4′,6-diamino-2-phenylindole (DAPI, 2µg /ml) for DNA staining or with 0.1% (v/v) SR2200 (Renchem) for callose staining and imaged using a LSM780 confocal microscope (Zeiss). Pollen count and viability were assessed using Alexander staining, described previously ^59^, and imaged with a Axioscope.A1 transmitted light microscope (Zeiss, 20×/0.5 objective), equipped with an Axiocam 105 camera and Visiview software (Visitron Systems). For GUS staining, plant tissues were fixed in 4% formaldehyde in 1X PBS, and washed in 0.1 M sodium phosphate buffer (pH 7.0). Tissues were infiltrated with GUS staining buffer (100 mM sodium phosphate buffer; pH-7.0, 10 mM EDTA; pH-8.0, 0.1% Triton X-100, 1 mM X-Gluc in DMF; 0.5 mM K_3_Fe(CN)_6_; 0.5 mM K_4_Fe(CN)_6_), by applying vacuum 15 minutes and incubated in dark at 37 °C for 48 hours. Stained tissues were imaged using a Stereo Discovery.V8 microscope (Zeiss). For cytological analysis of male meiosis, young inflorescences were collected from cultivated plants and fixed in ethanol:acetic acid (3:1, v/v) for 24 hours. Fixed tissues were rinsed twice in distilled water and twice in 10⍰mM sodium citrate buffer (pH 4.8), each for 5 minutes. Samples were then digested in citrate buffer containing 0.3% cellulase, cytohelicase, and pectolyase (Sigma-Aldrich) at 37⍰°C for 3 hours. Post-digestion, individual anthers were dissected onto slides in 20⍰μl glacial acetic acid, spread, and briefly heated on a 50⍰°C metal hot plate (∼30 seconds). Preparations were refixed in ethanol:acetic acid (3:1), air-dried with a hair dryer, and examined under phase contrast microscopy. Chromosomes were visualized using DAPI (2⍰μg/ml in Vectashield; Vector Laboratories) and imaged with a Zeiss Z2 epifluorescence microscope equipped with a CoolCube CCD camera.

### Immunocytology

For tubulin immunodetection, anthers were fixed in 4% formaldehyde in MTSB buffer (50 mM PIPES, 5 mM EGTA, 5 mM MgSO_4_, pH 6.9, 0.1% Triton X-100), dissected and gently squashed to release meiocytes onto glass slides using a pencil tip. The slides were snap-frozen in liquid nitrogen, and coverslips were removed with a razor blade. After air-drying at room temperature for 1 h, slides were rehydrated with a drop of MTSB for 5 min. Each slide was treated with 50 µL of 2% Driselase (Sigma-Aldrich, cat. no. D9515) in 7% (w/v) sucrose prepared in MTSB, covered with parafilm, and incubated in a humid chamber at room temperature for 30 min. Following enzymatic digestion, slides were washed twice with MTSB (5 min each) and incubated overnight at 4 °C with 150 µL of rat monoclonal anti-α-tubulin antibody (Serotec, cat. no. MCA78G) diluted 1:1000 in MTSB. The following day, slides were washed twice with MTSB (5 min each) and incubated for 1 h at room temperature with Cy3-conjugated goat anti-rat IgG secondary antibody (Chemicon, cat. no. AP183C) diluted 1:1000 in MTSB. After three washes with 200 µL of MTSB, slides were mounted in Vectashield containing DAPI and imaged using a LSM780 confocal microscope (Zeiss).

### Live cell imaging

Live-cell imaging of Arabidopsis pollen mother cells was performed using light sheet microscopy, as previously described ^4^. Anthers were imaged with a Light Sheet Z.1 microscope (Zeiss) equipped with a W Plan-Apochromat 20×/1.0 DIC objective, at 5-minute intervals.

### Protein localization in pollen mother cells

Inflorescences were fixed in 4% formaldehyde in 1X PBS for 15 min under vacuum, followed by 45 min of incubation without vacuum. The samples were washed three times for 5min with 1X PBS. Meiotic floral buds were then dissected to isolate meiotic anthers, which were incubated for 1 hour in 1X PBS containing 5 µg/ml DAPI. Subsequently, the anthers were washed twice with 1X PBS and then incubated at 60 °C in 200 µl of 1X PBS for 10 min in the dark, followed by a 10 min incubation on ice. A final wash was performed with 1 ml of 1X PBS. The anthers were then collected by centrifugation at 400 X g for 30 s, and the supernatant was carefully removed by pipetting, leaving approximately 20-40 µl of PBS. For imaging, the anthers were gently mounted on microscope slides using a cut 100 µl pipette tip to avoid mechanical damage, mixed with 20 µl of VECTASHIELD® Antifade Mounting Medium, and imaged using a Zeiss LSM780 confocal microscope equipped with a C-Apochromat 63×/1.2 W Korr UV-VIS-IR M27 water-immersion objective. Super-resolution microscopy was performed with Elyra 7 microscope (Zeiss) using lattice illumination pattern for 3D structured illumination (Lattice SIM), a Plan-Apochromat 63x/1.4 Oil DIC M27 objective, and dual PCO edge sCMOS cameras (1280 x 1280, pixel size 6.5 μm × 6.5 μm). Excitation was achieved with 488 nm and 561nm lasers. Raw images were acquired using at 1024 x 1024 pixels (pixel size 63 nm) with 9 phases. 2D image analyses were performed using Fiji ^60^. Depending on the experiment, either single optical slices or maximum intensity projections (MIPs) of five central slices were used for representative images and quantification, as specified in the figure legends. 3D rendering of RNP granules was carried out in Imaris v10.2.0 (Oxford Instruments) using volume rendering.

### Protein localization in protoplasts

Plasmids for protoplast transfection were prepared as follows. The coding-sequence of CDM1 (without introns and stop codon) was obtained by PCR amplification using primers CDM1-F and CDM1-RSTOP− with cDNA from floral tissue as the template. The PCR product was cloned into pENTR™/D-TOPO™, and the resulting entry clone was used in LR reactions with the BiFC-compatible destination vectors pGWnY and pGWcY ^57^, generating the constructs *35Sp::CDM1CDS:nYFP* and *35Sp::CDM1CDS:cYFP*, respectively. The same CDM1 entry clone was used in an LR reaction with the destination vector pGWB651 to generate the *35Sp::CDM1:G3GFP* construct for expression in protoplasts. Arabidopsis mesophyll protoplasts were isolated and transfected as previously described ^21, 61^ and imaged using LSM 780 confocal microscope (Zeiss) equipped with LCI Plan-Neofluar 63x/1.3 Imm Korr DIC M27 objective.

### FRAP analysis

Fluorescence recovery after photobleaching (FRAP) was performed using a ZEISS LSM780 confocal microscope. Regions of interest (ROIs) in protoplasts and meiocytes were bleached using a 514⍰nm laser at 100% intensity to deplete the GFP signal. Recovery of fluorescence was monitored by time-lapse imaging over 180 seconds. Image processing and quantification were conducted on maximum intensity projections (MIPs) of Z-stacks using Fiji.

### Single-molecule RNA Fluorescence In Situ Hybridization (smFISH)

smFISH was performed as previously described with some modifications ^62, 63^. Inflorescences were fixed in 4% methanol-free formaldehyde (Thermo Scientific, cat. no. 28908) in 1X PBST (10X PBS, Invitrogen, cat. no. AM9624, diluted in nuclease-free water [BioConcept, cat. no. 3-07F04-I] with 0.1% Triton X-100) under vacuum for 15 min, followed by 45 min of incubation without vacuum. Meiotic anthers were dissected under a stereo microscope, transferred onto a glass slide and covered with a coverslip. Excess buffer was removed by inverting the slide onto tissue paper. Anthers were gently squashed using a pencil tip to release individual meiocytes. Slides were immediately snap-frozen in liquid nitrogen, the coverslip removed with a razor blade, and slides were air-dried at room temperature for 1 hour. Slides were hydrated with 150⍰µL of pre-hybridization buffer (Stellaris Buffer A 1X [Biosearch Technologies, cat. no. SMF-WA1-60] containing 20% [v/v] formamide (AppliChem, cat. no. A2156) in nuclease-free water) for 5 minutes. Hybridization was carried out overnight at 37⍰°C in a dark, humid chamber using 150⍰µL of hybridization buffer (Stellaris Hybridization Buffer [Biosearch Technologies, cat. no. SMF-HB1-10] with 10% [v/v] formamide) containing 250 nM smFISH probe (Table S4). The samples were covered with a stripe of plastic foil and incubated in a dark humid chamber at 37^0^C for overnight. Slides were washed twice with pre-hybridization buffer at 37⍰°C in a dark, humid chamber (5 min and 25 min), then stained with 150⍰µL of DAPI (2⍰mg/mL) diluted in pre-hybridization buffer for 30 minutes at 37⍰°C. Finally, slides were washed with 200⍰µL of Stellaris Wash Buffer B (Biosearch Technologies, cat. no. SMF-WB1-20) for 5 minutes, mounted in VECTASHIELD®, and sealed with nail polish.

Imaging was performed using an Elyra 7 microscope (Zeiss) in widefield mode with Z-stacks acquired at 0.2 µm intervals. For G3BP5 or BT3 probes labeled with Quasar 570, the imaging settings were: excitation at 561 nm, 10% laser power, 500 ms exposure, and emission at 590 nm. For the GAPDH probe labeled with Quasar 670, the settings were: excitation at 642 nm, 30% laser power, 500 ms exposure, and emission at 655 nm. For DAPI, excitation was at 405 nm, with 3% laser power, 100 ms exposure, and emission at 440 nm. Probe sequences and catalog numbers are listed in Table S2.

### RNA immuno-Precipitation (RIP)

Approximately 50-100⍰mg of 0.3 – 0.5 mm floral buds from *Arabidopsis* wild-type and *CDM1p::CDM1:G3GFP* plants were collected in a watch glass on ice. After removing surface moisture with a tissue towel, buds were snap-frozen in liquid nitrogen and ground into a fine powder using mortar and pestle. Lysis was initiated by adding 1⍰mL of ice-cold lysis buffer (50⍰mM Tris-HCl pH⍰7.5, 150⍰mM NaCl, 5⍰mM MgCl_2_, 0.5⍰mM DTT, 10% glycerol, 0.05% Nonidet P-40, 1⍰mM PMSF, 1X protease inhibitor cocktail [cOmplete ULTRA Tablets, EDTA-free; Roche, cat. no. 05892953001], 20⍰U/mL RNaseOUT [Invitrogen, cat. no. 10777019]) prepared in DEPC treated, sterile MilliQ water, directly into the mortar. The lysate was thawed to 4⍰°C and transferred to RNase-free Protein LoBind microcentrifuge tubes, followed by incubation on ice for 30 minutes with occasional mixing. Samples were centrifuged at 10,000⍰rpm for 10 minutes at 4⍰°C. Protein concentration was estimated using the Bradford assay, and 1 mg of protein extract was used for immunoprecipitation with ChromoTek GFP-Trap® magnetic beads.

To reduce nonspecific binding, lysates were pre-cleared before pull-down with 30ul of ChromoTek binding control beads (lacking antibody), and incubated with lysates on a rotating wheel at 4⍰°C for 45 min. Five percent of the pre-cleared lysates was reserved as input and stored at -20⍰°C. In parallel, GFP-Trap beads were equilibrated in lysis buffer. Pre-cleared lysates were incubated with 30 µL of equilibrated GFP-Trap beads for 90 min at 4⍰°C with rotation at 16rpm. Beads were collected using a magnetic rack and washed three times with wash buffer (50⍰mM Tris-HCl pH⍰7.5, 150⍰mM NaCl, 5⍰mM MgCl_2_, 0.5⍰mM DTT, 20⍰U/mL RNaseOUT) for 5 min each at 4°C. The final wash was transferred to new DNA LoBind tubes.

### RNA Preparation and NGS Library Preparation

RNA was extracted from both input and immunoprecipitated samples by adding 1 mL of RNA Blue reagent (Top-Bio, cat. no. R013) and processing according to the manufacturer’s instructions. RNA concentrations were measured using the Qubit RNA Assay Kit (Invitrogen, cat. no. Q10210), and residual DNA was removed with TURBO DNase (Invitrogen, cat. no. AM1907). For NGS library preparation, 1–5 ng of DNA-free RNA was used. Libraries were prepared using the TruSeq Stranded Total RNA Library Prep Kit (Illumina) and indexed with IDT for Illumina TruSeq DNA and RNA UD Indexes, following the manufacturer’s protocol. Sequencing was performed on an Illumina NextSeq 500 platform using the NextSeq 500/550 High Output Kit v2.5 (75 cycles).

### RIP-seq data analysis

Raw sequencing reads were quality-checked using FastQC v0.12.1 (https://www.bioinformatics.babraham.ac.uk/projects/fastqc/). Low-quality reads and adapter contaminations were removed with cutadapt v3.5 ^64^ . The high-quality reads were then aligned to the Arabidopsis thaliana reference transcriptome AtRTD2 dataset ^65^ and simultaneously quantified with mapping-based algorithm using Salmon v10 ^66^ . The downstream data normalization and differential expression analysis was done using DESeq2 ^67^ in R (v4.4.2). The log2 fold change (log2FC) for each gene was estimated using the Wald test. Due to the lack of biological replicates, gene-wise dispersions could not be estimated empirically. Therefore, we specified a fixed dispersion value of 0.1 for all genes, which corresponds to a moderate level of variability commonly observed in RNA-seq datasets. While this approach enables statistical testing, the resulting p-values should be interpreted with caution and genes with high log2FC (>2) and adequate p-values (p-value < 0.05) were considered as putative candidates for further validation.

### cDNA Preparation and qPCR Analysis

cDNA was synthesized from 500⍰ng of DNA-free RNA from floral buds or RNA prepared after RIP, using the SuperScript™ IV (Invitrogen, cat. no. 18090050), following the manufacturer’s protocol. A mix of random hexamer primers (Thermo Scientific, cat. no. SO142) and oligo(dT) primers (Thermo Scientific, cat. no. SO132) were used for reverse transcription. Quantitative PCR (qPCR) was performed using the KAPA SYBR® FAST qPCR Kit (KAPA Biosystems, cat. no. KR0389). Primer sequences used for qPCR are listed in Table S3.

### Protein expression and purification

For bacterial expression, CDM1 cDNA was amplified from the 35Sp::CDM1:G3GFP template using either CDM1-F and G3GFP-R primers to generate the CDM1-G3GFP fragment, or the CDM1-F and CDM1-RSTOP primers to generate the CDM1 fragment. The PCR products were cloned into the SspI-linearized pET His6-MBP-TEV LIC vector (1M). The resulting constructs, *His-MBP-CDM1-G3GFP* and *His-MBP-CDM1*, were transformed into E. coli BL21-Rosetta cells for protein expression in LB medium containing kanamycin. Protein expression was induced with IPTG (0.5⍰mM) for 4 hours at 30⍰°C when the culture reached an OD_600_ of 0.4. Cells were harvested by centrifugation, and the pellet was resuspended in lysis buffer containing 0.1⍰M Tris-HCl (pH⍰7.5), 0.5⍰M NaCl, 0.05% Tween 20, 1% Triton X-100, 1⍰mg/mL lysozyme (Sigma-Aldrich, cat. no. L6876), 1⍰mM DTT, and 1X protease inhibitor cocktail (cOmplete ULTRA Tablets, EDTA-free; Roche, cat. no. 05892953001). The suspension was sonicated and centrifuged, and the resulting supernatant was filtered through a 0.2⍰µm filter (Sartorius, cat. no. 17764) and supplemented with 50⍰mM imidazole (Sigma-Aldrich, cat. no. I5513). His-tagged proteins were purified using His Mag Sepharose™ Ni magnetic beads (Cytiva, cat. no. 28967390) following the manufacturer’s protocol. Eluted His-MBP-CDM1-GFP was dialyzed into storage buffer (20⍰mM HEPES, pH⍰7.4, 150⍰mM KCl, 1⍰mM DTT), while His-MBP-CDM1 was dialyzed in RNase free 1X PBS, concentrated using Amicon® Ultra-0.5 centrifugal filters (Merck Millipore, cat. no. UFC503024). Purified proteins were stored at −80⍰°C.

### In vitro dimerization assay and Western blot

20⍰µg of purified His_6_-MBP-CDM1-GFP protein was incubated with His_6_-tagged TEV protease (Promega, cat. no. V610A) in 100⍰µL total volume (1X TEV buffer, 1⍰mM DTT, 2⍰µL TEV) at room temperature for 1 hour. The reaction was incubated with 10⍰µL His Mag Sepharose™ Ni beads for 30 minutes at room temperature, followed by a second round of purification with fresh beads. The cleaved unbound CDM1-GFP was collected for further analysis. For the dimerization assay, equimolar amounts of His_6_-MBP-CDM1-GFP and CDM1-GFP were mixed in storage buffer (20⍰mM HEPES, pH⍰7.4, 150⍰mM KCl, 1⍰mM DTT) in a 50⍰µL reaction and incubated at room temperature for 2 hours on a rotating wheel (16⍰rpm). Complexes were immunoprecipitated by incubating with 1⍰µL anti-His antibody (Abcam, cat. no. ab18184) for 30min, followed by Protein G-conjugated Sepharose beads (Invitrogen, cat. no. 10003D) pre-blocked with 2% BSA for 30 min at room temperature. Beads were washed with storage buffer, resuspended in 20⍰µL, and stored at -20⍰°C along with input and unbound fractions. Proteins were eluted with 1X Laemmli buffer and boiled prior to SDS-PAGE. Western blots were probed with rat anti-GFP antibody (Chromotek, cat. no. 3H9) and HRP-conjugated secondary antibody. Detection was performed using a luminol-based chemiluminescent substrate (Thermo Scientific, cat. no. 34095). Imaging was performed using ChemiDoc XRS+ system (BioRad).

### In vitro phase separation assay

Phase separation assays were performed by mixing increasing concentrations of purified His_6_-MBP-CDM1-GFP in buffer (20⍰mM HEPES, pH⍰7.4, 150⍰mM KCl, 1⍰mM DTT). To remove the N-terminal solubilizing His-MBP tag, 0.5⍰U/µL TEV protease was added, and phase separation was induced by adding 10% (w/v) PEG8000 (Carl Roth, cat. no. 0263.2). The effect of RNA was tested by adding 100⍰ng/µL *in vitro* transcribed 5⍰ UTR RNA of *G3BP5*. 5⍰µL of the mixture was transferred into custom-made chambers constructed using double-sided tape on dichlorodimethylsilane-treated microscopy slides. Chambers were sealed with a coverslip, inverted, and incubated in a humidified chamber at 25⍰°C for 10 minutes. Droplet formation was visualized using a ZEISS LSM780 confocal microscope with a C-Apochromat 63×/1.2 W Korr UV-VIS-IR M27 objective.

### RNA Electrophoretic Mobility Shift Assay (RNA-EMSA)

RNA probes covering the three distinct regions at the 5′end of *G3BP5* mRNA were prepared by *in vitro* transcription of DNA templates amplified by PCR using the following primer pairs: T7-1F and 75R, T7-75F and 162R, and T7-112F and 188R (Table S3). The PCR products were purified from the agarose gels using the NucleoSpin® Gel and PCR Clean-Up Kit (Macherey-Nagel), extracted once with phenol:chloroform:IAA (25:24:1), twice with chloroform:IAA (24:1), ethanol-precipitated, washed with 75% RNase-free ethanol, and re-suspended in 10 µL of DEPC-treated water. RNA concentrations were determined using a NanoDrop spectrophotometer. Radiolabeled RNA probes and unlabeled competitor RNA were synthesized by *in vitro* transcription from 1 µg of the DNA template using the HiScribe T7 High Yield RNA Synthesis Kit (New England Biolabs) according to manufacturer′s instructions. For the radiolabeled transcripts, α-^32^P-UTP (3 µL, 10 mCi/mL, >6000 Ci/mmol) was included in the reaction. After transcription, DNA templates were removed using the TURBO DNA-free kit (Thermo Scientific), and RNA was purified by RNA-blue reagent extraction (Top-Bio) followed by two chloroform:IAA extractions and ethanol precipitation. The RNA was resuspended in RNase-free PBS buffer, matching the dialysis buffer of the His-MBP-CDM1 protein. RNA concentrations were quantified using Qubit 3.0 Fluorometer (Thermo Fisher).

Binding reactions (final volume of 20 µL) contained 20 nM of radiolabeled RNA, increasing concentrations of His-MBP-CDM1 protein, 10 mM HEPES, pH 7.3, 20 mM KCl, 1 mM MgCl_2_, 10 µM ZnSO_4_, 1 mM DTT, 5% glycerol, 0.1 mg/mL heparin, and 0.5 U of ProTEV Plus protease (Promega). Reactions were incubated at 37 °C for 15 minutes and then resolved on native 5% polyacrylamide gels (acrylamide:bisacrylamide, 29:1; Sigma) prepared in 0.5× TBE. Gels were pre-electrophoresed in 0.5× TBE at 100 V for 1 h at 4 °C, run with samples at 160 V for 5 h at 4 °C, dried, exposed to phosphor screens (GE Healthcare), and scanned using a Typhoon™ FLA 7000 imaging system (GE Healthcare).

### Yeast Two-Hybrid Assay

Yeast two-hybrid experiments were performed using the Matchmaker™ GAL4-based system (Clontech). Bait and prey constructs were generated according to the manufacturer’s instructions using cDNA synthesized from RNA isolated from floral tissue and transformed into *Saccharomyces cerevisiae* strains Y187 and AH109, respectively. Protein-protein interactions were assayed by the mating method. Diploid yeast cells containing both plasmids were selected on SD/-Leu/-Trp medium and screened for interactions on SD/-Leu/-Trp/-His medium supplemented with 2.5 mM 3-amino-1,2,4-triazole (3-AT) at 30 °C. Empty vector combinations were included as negative controls to test for autoactivation.

## Acknowledgements

This work was supported by the Czech Science Foundation (EXPRO grant 23-07969X). We further acknowledge the core facilities of CEITEC Masaryk University for support in obtaining and analyzing scientific data presented in this paper: CELLIM funded by MEYS CR (LM2023050 Czech-BioImaging), Plant Science Core Facility, and the CF Genomics supported by the NCMG research infrastructure (LM2023067 funded by MEYS CR).

---

**Video 1**. Live imaging of meiosis in pollen mother cells expressing CDM1-GFP. Scale bar = 20 µm.

## References

1. Bennett, M.D. The time and duration of meiosis. Philos Trans R Soc Lond B Biol Sci 277, 201–226 (1977).

2. Nasar, S.K.T. & Rai, R. Seasonal Fluctuation of the Duration of Meiosis in Rhoeo. Environ Exp Bot 25, 253–256 (1985).

3. Prusicki, M.A. et al. Live cell imaging of meiosis in Arabidopsis thaliana. Elife 8 (2019).

4. Valuchova, S. et al. Imaging plant germline differentiation within Arabidopsis flowers by light sheet microscopy. Elife 9 (2020).

5. Kolowerzo-Lubnau, A., Niedojadlo, J., Swidzinski, M., Bednarska-Kozakiewicz, E. & Smolinski, D.J. Transcriptional activity in diplotene larch microsporocytes, with emphasis on the diffuse stage. PLoS One 10, e0117337 (2015).

6. Charalambous, C., Webster, A. & Schuh, M. Aneuploidy in mammalian oocytes and the impact of maternal ageing. Nat Rev Mol Cell Biol 24, 27–44 (2023).

7. Alexander, A.K. et al. A-MYB and BRDT-dependent RNA Polymerase II pause release orchestrates transcriptional regulation in mammalian meiosis. Nat Commun 14, 1753 (2023).

8. Bellutti, L. et al. Genome-wide transcriptional silencing and mRNA stabilization allow the coordinated expression of the meiotic program in mice. Nucleic Acids Res 53 (2025).

9. Gao, J., Qin, Y. & Schimenti, J.C. Gene regulation during meiosis. Trends Genet 40, 326–336 (2024).

10. Jin, L. & Neiman, A.M. Post-transcriptional regulation in budding yeast meiosis. Curr Genet 62, 313–315 (2016).

11. Morgan, M., Kumar, L., Li, Y. & Baptissart, M. Post-transcriptional regulation in spermatogenesis: all RNA pathways lead to healthy sperm. Cell Mol Life Sci 78, 8049–8071 (2021).

12. Brar, G.A. et al. High-resolution view of the yeast meiotic program revealed by ribosome profiling. Science 335, 552–557 (2012).

13. Carlile, T.M. & Amon, A. Meiosis I is established through division-specific translational control of a cyclin. Cell 133, 280–291 (2008).

14. Berchowitz, L.E. et al. Regulated Formation of an Amyloid-like Translational Repressor Governs Gametogenesis. Cell 163, 406–418 (2015).

15. Jin, L., Zhang, K., Xu, Y., Sternglanz, R. & Neiman, A.M. Sequestration of mRNAs Modulates the Timing of Translation during Meiosis in Budding Yeast. Mol Cell Biol 35, 3448–3458 (2015).

16. da Cruz, I. et al. Transcriptome analysis of highly purified mouse spermatogenic cell populations: gene expression signatures switch from meiotic-to postmeiotic-related processes at pachytene stage. BMC Genomics 17, 294 (2016).

17. Susor, A., Jansova, D., Anger, M. & Kubelka, M. Translation in the mammalian oocyte in space and time. Cell Tissue Res 363, 69–84 (2016).

18. Cheng, S. et al. Mammalian oocytes store mRNAs in a mitochondria-associated membraneless compartment. Science 378, eabq4835 (2022).

19. Rudzka, M. et al. Functional nuclear retention of pre-mRNA involving Cajal bodies during meiotic prophase in European larch (Larix decidua). Plant Cell 34, 2404–2423 (2022).

20. Cairo, A. et al. SMG7 and eIF4A constitute a homeostatic module controlling P-body condensation and function of Meiotic bodies. bioRxiv, 2025.2005.2003.652043 (2025).

21. Cairo, A. et al. Meiotic exit in Arabidopsis is driven by P-body-mediated inhibition of translation. Science 377, 629–634 (2022).

22. Bulankova, P., Riehs-Kearnan, N., Nowack, M.K., Schnittger, A. & Riha, K. Meiotic progression in Arabidopsis is governed by complex regulatory interactions between SMG7, TDM1, and the meiosis I-specific cyclin TAM. Plant Cell 22, 3791–3803 (2010).

23. Cifuentes, M. et al. TDM1 Regulation Determines the Number of Meiotic Divisions. PLoS Genet 12, e1005856 (2016).

24. d’Erfurth, I. et al. The cyclin-A CYCA1;2/TAM is required for the meiosis I to meiosis II transition and cooperates with OSD1 for the prophase to first meiotic division transition. PLoS Genet 6, e1000989 (2010).

25. Nelms, B. & Walbot, V. Gametophyte genome activation occurs at pollen mitosis I in maize. Science 375, 424–429 (2022).

26. Pomeranz, M.C. et al. The Arabidopsis tandem zinc finger protein AtTZF1 traffics between the nucleus and cytoplasmic foci and binds both DNA and RNA. Plant Physiol 152, 151–165 (2010).

27. Lu, P. et al. The Arabidopsis CALLOSE DEFECTIVE MICROSPORE1 gene is required for male fertility through regulating callose metabolism during microsporogenesis. Plant Physiol 164, 1893–1904 (2014).

28. Chai, G. et al. Arabidopsis C3H14 and C3H15 have overlapping roles in the regulation of secondary wall thickening and anther development. J Exp Bot 66, 2595–2609 (2015).

29. Capitao, C. et al. A CENH3 mutation promotes meiotic exit and restores fertility in SMG7-deficient Arabidopsis. PLoS Genet 17, e1009779 (2021).

30. Riehs-Kearnan, N., Gloggnitzer, J., Dekrout, B., Jonak, C. & Riha, K. Aberrant growth and lethality of Arabidopsis deficient in nonsense-mediated RNA decay factors is caused by autoimmune-like response. Nucleic Acids Res 40, 5615–5624 (2012).

31. De Storme, N. & Geelen, D. Cytokinesis in plant male meiosis. Plant Signal Behav 8, e23394 (2013).

32. Kim, W.C. et al. AtC3H14, a plant-specific tandem CCCH zinc-finger protein, binds to its target mRNAs in a sequence-specific manner and affects cell elongation in Arabidopsis thaliana. Plant J 80, 772–784 (2014).

33. Klepikova, A.V., Kasianov, A.S., Gerasimov, E.S., Logacheva, M.D. & Penin, A.A. A high resolution map of the Arabidopsis thaliana developmental transcriptome based on RNA-seq profiling. Plant J 88, 1058–1070 (2016).

34. Hardy, E.C. & Balcerowicz, M. Untranslated yet indispensable-UTRs act as key regulators in the environmental control of gene expression. J Exp Bot 75, 4314–4331 (2024).

35. Laureau, R. et al. Meiotic Cells Counteract Programmed Retrotransposon Activation via RNA-Binding Translational Repressor Assemblies. Dev Cell 56, 22–35 e27 (2021).

36. Rojas, J. et al. Spo13/MEIKIN ensures a Two-Division meiosis by preventing the activation of APC/C(Ama1) at meiosis I. EMBO J 42, e114288 (2023).

37. Brangwynne, C.P. et al. Germline P granules are liquid droplets that localize by controlled dissolution/condensation. Science 324, 1729–1732 (2009).

38. Pamula, M.C. & Lehmann, R. How germ granules promote germ cell fate. Nat Rev Genet 25, 803–821 (2024).

39. Cassani, M. & Seydoux, G. P-body-like condensates in the germline. Semin Cell Dev Biol 157, 24–32 (2024).

40. Cardona, A.H. et al. Self-demixing of mRNA copies buffers mRNA:mRNA and mRNA:regulator stoichiometries. Cell 186, 4310–4324 e4323 (2023).

41. Komiya, R. Spatiotemporal regulation and roles of reproductive phasiRNAs in plants. Genes Genet Syst 96, 209–215 (2022).

42. Cheng, J. & Martinez, G. Enjoy the silence: Canonical and non-canonical RNA silencing activity during plant sexual reproduction. Curr Opin Plant Biol 82, 102654 (2024).

43. Bogamuwa, S.P. & Jang, J.C. Tandem CCCH zinc finger proteins in plant growth, development and stress response. Plant Cell Physiol 55, 1367–1375 (2014).

44. Pomeranz, M., Lin, P.C., Finer, J. & Jang, J.C. AtTZF gene family localizes to cytoplasmic foci. Plant Signal Behav 5, 190–192 (2010).

45. Maldonado-Bonilla, L.D. et al. The Arabidopsis tandem zinc finger 9 protein binds RNA and mediates pathogen-associated molecular pattern-triggered immune responses. Plant Cell Physiol 55, 412–425 (2014).

46. Guo, C. et al. RNA Binding Protein OsTZF7 Traffics Between the Nucleus and Processing Bodies/Stress Granules and Positively Regulates Drought Stress in Rice. Front Plant Sci 13, 802337 (2022).

47. Qu, J., Kang, S.G., Wang, W., Musier-Forsyth, K. & Jang, J.C. The Arabidopsis thaliana tandem zinc finger 1 (AtTZF1) protein in RNA binding and decay. Plant J 78, 452–467 (2014).

48. He, S.L. et al. Overexpression of stress granule protein TZF1 enhances salt stress tolerance by targeting ACA11 mRNA for degradation in Arabidopsis. Front Plant Sci 15, 1375478 (2024).

49. Chai, G. et al. Integration of C3H15-mediated transcriptional and post-transcriptional regulation confers plant thermotolerance in Arabidopsis. Plant J 119, 1558–1569 (2024).

50. Robert, H.S., Quint, A., Brand, D., Vivian-Smith, A. & Offringa, R. BTB and TAZ domain scaffold proteins perform a crucial function in Arabidopsis development. Plant J 58, 109–121 (2009).

51. Pelechano, V., Wei, W. & Steinmetz, L.M. Widespread Co-translational RNA Decay Reveals Ribosome Dynamics. Cell 161, 1400–1412 (2015).

52. Yu, X., Willmann, M.R., Anderson, S.J. & Gregory, B.D. Genome-Wide Mapping of Uncapped and Cleaved Transcripts Reveals a Role for the Nuclear mRNA Cap-Binding Complex in Cotranslational RNA Decay in Arabidopsis. Plant Cell 28, 2385–2397 (2016).

53. Deragon, J.M. & Merret, R. Co-Translational mRNA Decay in Plants: Recent advances and future directions. J Exp Bot (2025).

54. Hubstenberger, A., Noble, S.L., Cameron, C. & Evans, T.C. Translation repressors, an RNA helicase, and developmental cues control RNP phase transitions during early development. Dev Cell 27, 161–173 (2013).

55. De Jaeger-Braet, J. et al. The recruitment of the A-type cyclin TAM to stress granules is crucial for meiotic fidelity under heat. Sci Adv 11, eadr5694 (2025).

56. Enami, K. et al. Differential expression control and polarized distribution of plasma membrane-resident SYP1 SNAREs in Arabidopsis thaliana. Plant Cell Physiol 50, 280–289 (2009).

57. Nakagawa, T. et al. Improved Gateway binary vectors: high-performance vectors for creation of fusion constructs in transgenic analysis of plants. Biosci Biotechnol Biochem 71, 2095–2100 (2007).

58. Javorka, P., Raxwal, V.K., Najvarek, J. & Riha, K. artMAP: A user-friendly tool for mapping ethyl methanesulfonate-induced mutations in Arabidopsis. Plant Direct 3, e00146 (2019).

59. Alexander, M.P. Differential staining of aborted and nonaborted pollen. Stain Technol 44, 117–122 (1969).

60. Schindelin, J. et al. Fiji: an open-source platform for biological-image analysis. Nat Methods 9, 676–682 (2012).

61. Yoo, S.D., Cho, Y.H. & Sheen, J. Arabidopsis mesophyll protoplasts: a versatile cell system for transient gene expression analysis. Nat Protoc 2, 1565–1572 (2007).

62. Duncan, S., Olsson, T.S.G., Hartley, M., Dean, C. & Rosa, S. A method for detecting single mRNA molecules in Arabidopsis thaliana. Plant Methods 12, 13 (2016).

63. Duncan, S., Johansson, H.E. & Ding, Y. Reference genes for quantitative Arabidopsis single molecule RNA fluorescence in situ hybridization. J Exp Bot 74, 2405–2415 (2023).

64. Martin, M. Cutadapt removes adapter sequences from high-throughput sequencing reads. EMBnet.journal; Vol 17, No 1: Next Generation Sequencing Data Analysis DO - 10.14806/ej.17.1.200 (2011).

65. Zhang, R. et al. A high quality Arabidopsis transcriptome for accurate transcript-level analysis of alternative splicing. Nucleic Acids Res 45, 5061–5073 (2017).

66. Patro, R., Duggal, G., Love, M.I., Irizarry, R.A. & Kingsford, C. Salmon provides fast and bias-aware quantification of transcript expression. Nat Methods 14, 417–419 (2017).

67. Love, M.I., Huber, W. & Anders, S. Moderated estimation of fold change and dispersion for RNA-seq data with DESeq2. Genome Biol 15, 550 (2014).

