## Supplementary Figures S1-S7 for "Post-transcriptional regulation of meiotic transcripts by the RNA binding protein CDM1 is associated with cytoplasmic condensate assemblies"

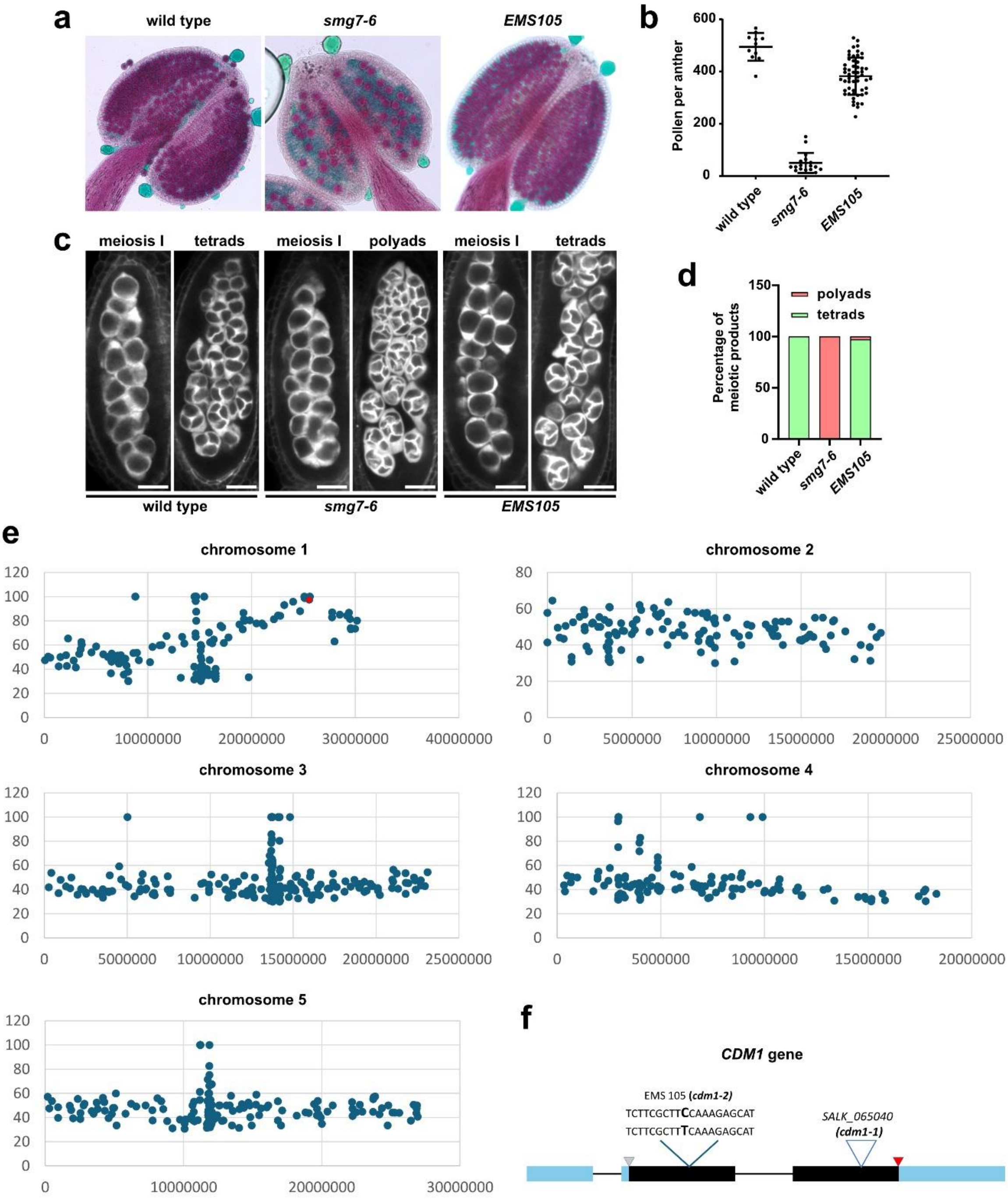


**Figure S1. Identification of *cdm1-2* allele in a forward genetic screen.**

(**a, b**) A suppressor screen for mutants with increased fertility in the *smg7-6* background led to the identification of the recessive EMS105 line, which showed restored pollen production. (**a**) Alexander staining of wild type, *smg7-6*, and EMS105 pollen. (**b**) Quantification of pollen viability in the EMS105 line.

(**c, d**) *smg7-6* mutants form polyads due to aberrant meiotic exit and unequal chromosome segregation at the end of meiosis. EMS105 suppresses this defect and restores the formation of regular tetrads.

(**c**) SR2200 staining of callose during meiosis I and at the end of meiosis. Scale bar: 20 µm. (**d**) Quantification of tetrads and polyads formed at the end of meiosis (n = 578, 316, and 670 for wild type, *smg7-6* and *EMS105,* respectively).

(**e**) Identification of *de novo* mutations associated with increased fertility in the segregating population generated by backcrossing EMS105 to *smg7-6*. Dot plots show the distribution and frequency of *de novo* mutations identified in fertile plants along individual chromosomes. The red dot on chromosome 1 marks a *de novo* mutation in *CDM1*. The individual mutations are presented in Table S1.

(**f**) Schematic representation of the *CDM1* gene, showing the positions of exons and introns, coding sequence (black), and the *cdm1-1* and *cdm1-2* mutations. The *cdm1-2* allele represents a C-to-T transition that causes a proline-to-serine substitution at position 152.

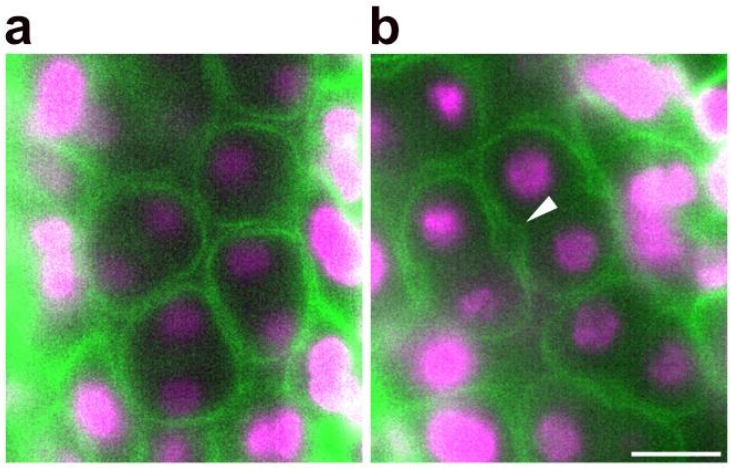


**Figure S2. Premature cytokinesis initiation in *cdm1-1* visualized using a plasma membrane marker.** (**a**) Light-sheet micrograph of a wild-type interkinesis meiocyte with nuclei labeled by the chromatin marker HTA10–RFP (magenta) and plasma membranes labeled by SYP132–GFP (green). The plasma membrane encircling the meiocyte shows no invaginations. (**b**) Interkinesis *cdm1-1* meiocyte showing premature plasma membrane invaginations, indicative of early cytokinesis initiation. The arrow marks a membrane invagination site. Scale bar, 10 µm.

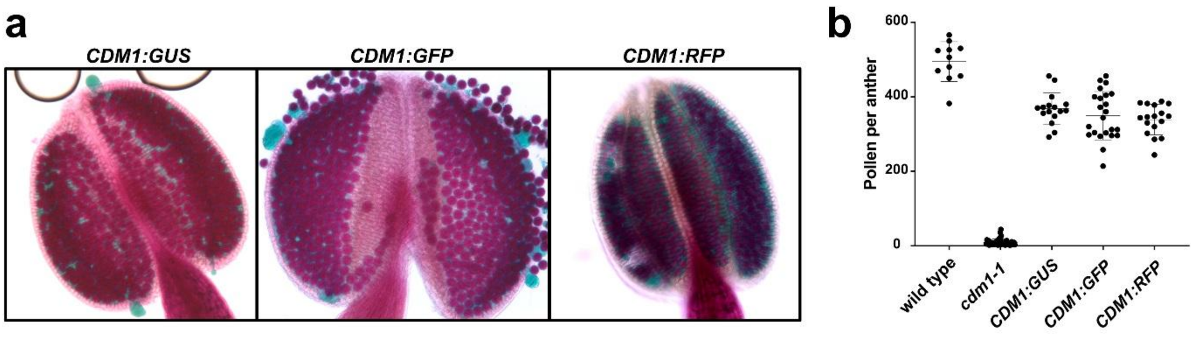


**Figure S3. Complementation of the *cdm1-1* with reporter lines.** (**a**) Alexander staining of wild type, *cdm1-1*, and complemented lines (*CDM1*:GUS, *CDM1*:GFP, *CDM1*:*RFP*). (**b**) Quantification of viable pollen showing restoration of pollen count in *cdm1-1* mutants complemented with the marker lines.

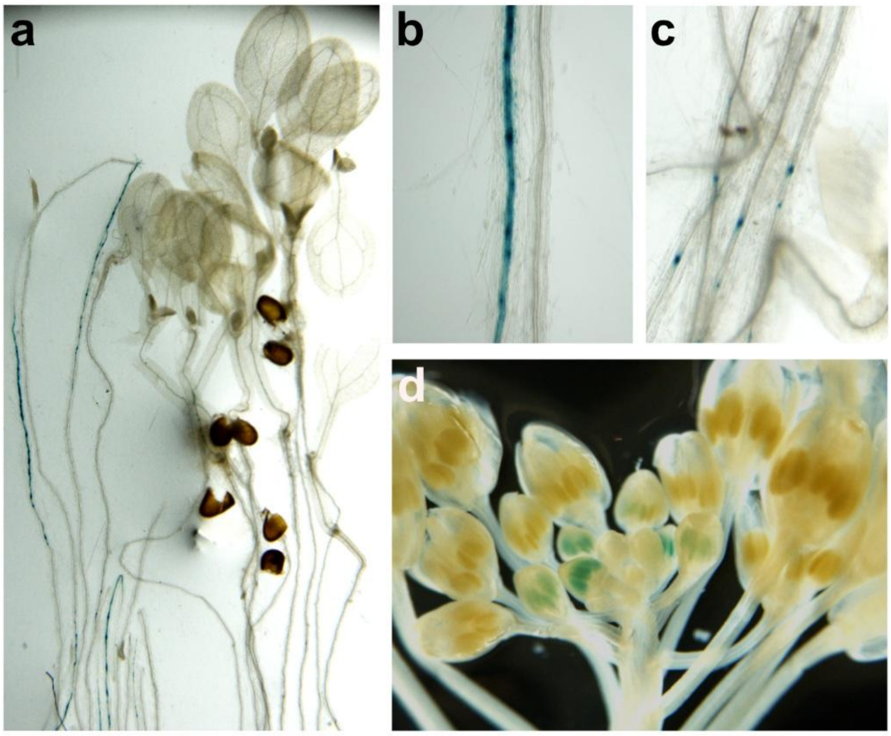


**Figure S4: Localization of *CDM1p*::*CDM1*:*GUS* expression.** (**a**) Histochemical GUS staining in 7-day-old seedlings shows punctate expression in the root vasculature, with no detectable signal in root tips, hypocotyls, or young leaves. (**b,c**) Detailed views of GUS staining in the root vasculature. (**d**) GUS staining in inflorescences of 4-week-old *CDM1p::CDM1:GUS* plants, showing strong expression in young meiotically active buds and no expression in older buds.

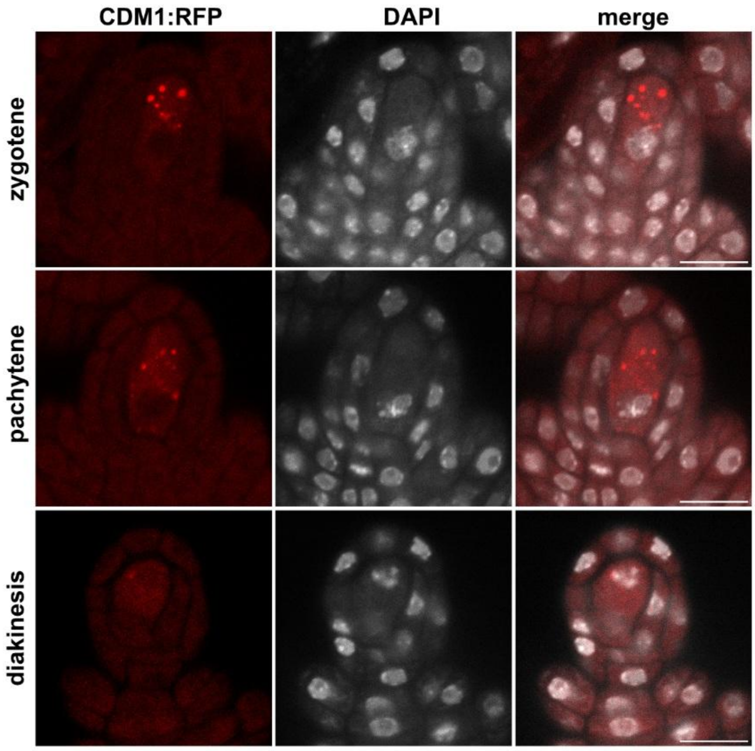


**Figure S5. Expression of CDM1:RFP in megaspore mother cells.** CDM1:RFP form cytoplasmic foci similar to those observed in pollen mother cells. DNA is counterstained with DAPI. Scale bar = 10 µm.

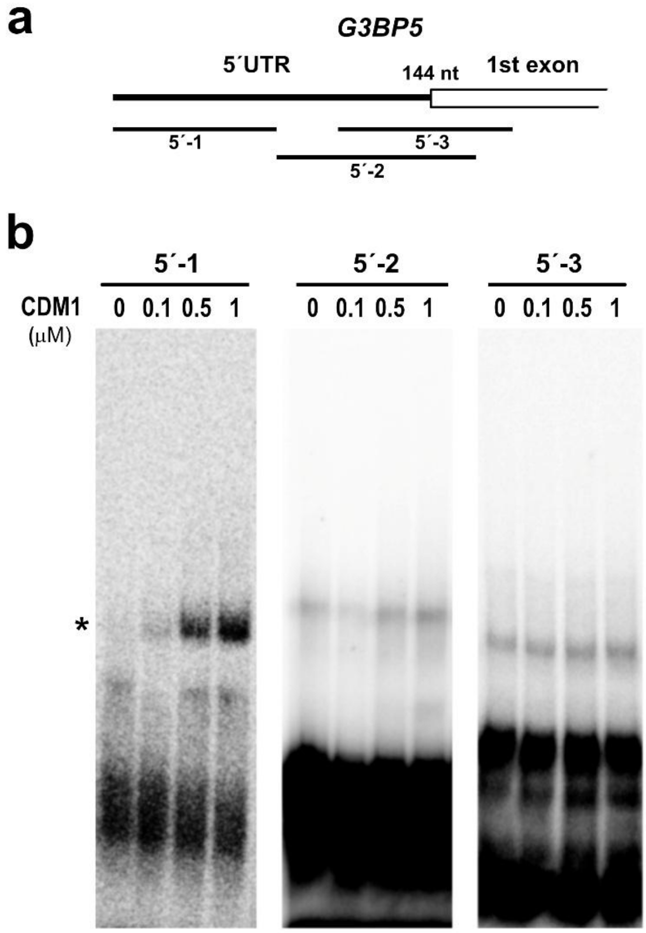


**Figure S6. Identification of *G3BP5* mRNA region interacting with CDM1 by REMSA.** (**a**) Schematic representation of the 5´ end of *G3BP5* mRNA, with the positions of probes used for REMSA indicated. (**b**) Autoradiogram of REMSA showing that only the 5′-1 RNA probe exhibits increased binding to CDM1 in a protein concentration-dependent manner (indicated by asterisk).

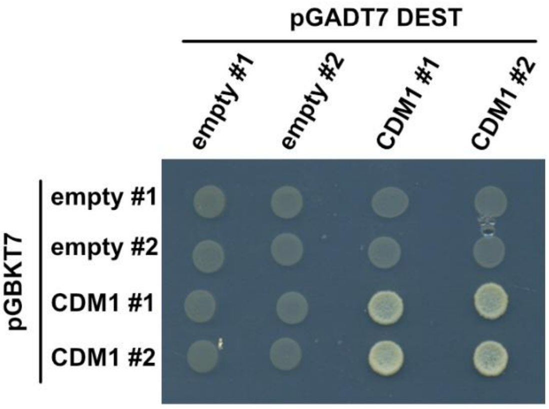


**Figure S7. CDM1 self-association detected by yeast two-hybrid assay.** Two independent clones were tested for each empty vector control and CDM1 construct.
